# Distribution of the gyromitrin mycotoxin in the lorchel family assessed by a pre-column-derivatization and ultra high-performance liquid chromatography method

**DOI:** 10.1101/2022.08.08.503034

**Authors:** Alden C. Dirks, Osama G. Mohamed, Pamela Schultz, Andrew N. Miller, Ashootosh Tripathi, Timothy Y. James

## Abstract

Gyromitrin (acetaldehyde *N*-methyl-*N*-formylhydrazone) and its homologs are deadly mycotoxins produced most infamously by the lorchel (also known as false morel) *Gyromitra esculenta*, which is paradoxically consumed as a delicacy in some parts of the world. There is much speculation about the presence of gyromitrin in other species of the lorchel family (Discinaceae), but no studies have broadly assessed its distribution. Given the history of poisonings associated with the consumption of *G. esculenta* and *G. ambigua*, we hypothesized that gyromitrin evolved in the last common ancestor of these taxa and would be present in their descendants with adaptive loss of function in the nested truffle clade, *Hydnotrya*. To investigate this hypothesis, we developed a sensitive analytical derivatization method for the detection of gyromitrin using 2,4-dinotrobenzaldehyde as the derivatization reagent. In total, we analyzed 66 specimens for the presence of gyromitrin over 105 tests. Moreover, we sequenced the nuc rDNA ITS1-5.8S-ITS2 region (ITS barcode) and nuc 28S rDNA to assist in species identification and to infer a supporting phylogenetic tree. We detected gyromitrin in all tested specimens from the *G. esculenta* group as well as *G. leucoxantha*. This distribution is consistent with a model of rapid evolution coupled with horizontal transfer, which is typical for secondary metabolites. We clarified that gyromitrin production in Discinaceae is both discontinuous and more limited than previously thought. Further research is required to elucidate the gyromitrin biosynthesis gene cluster and its evolutionary history in lorchels. KEYWORDS: 2,4-dinitrobenzaldehyde, *Gyromitra* spp., *Hydnotrya* spp., Discinaceae, Pezizales, Schiff bases, UHPLC-DAD analysis

## INTRODUCTION

Gyromitrin is a polar, water-soluble, and volatile mycotoxin produced by *Gyromitra esculenta*, a distinctive brain-like mushroom that is consumed as a delicacy, particularly in Scandinavia (Härkönen 1998; Svanberg and Lindh 2019; Benjamin 2020; Sitta et al. 2021). Structurally, gyromitrin refers to acetaldehyde *N*-methyl-*N*-formylhydrazone (**1**), and occurs with eight higher aldehyde homologs (**2–9**) present in smaller quantities (List and Luft 1968a; Pyysalo 1975; Pyysalo and Niskanen 1977) (FIG. 1). Spontaneously at room temperature, upon heating, and especially in acidic environments like the stomach, gyromitrin hydrolyzes into its aldehyde component and *N*-methyl-*N*-formylhydrazine (**10**), which then loses formaldehyde to yield monomethylhydrazine (**11**), an ingredient in some rocket propellants (List and Luft 1968b; Pyysalo et al. 1978; Monteith 2020) (FIG. 2). Gyromitrin’s volatility, solubility, and reactivity explain how *G. esculenta*, which contains 50–300 mg/kg of gyromitrin per fresh mushroom, can be consumed without ill effect (Pyysalo 1976; Pyysalo and Niskanen 1977; Michelot and Toth 1991). By boiling the mushrooms twice, rinsing them after each boil, and changing the cooking liquid in between, more than 99% of the gyromitrin is released to render the mushroom safe to eat without acute toxicity (Pyysalo 1976; Pyysalo and Niskanen 1977).

**Figure 1.**
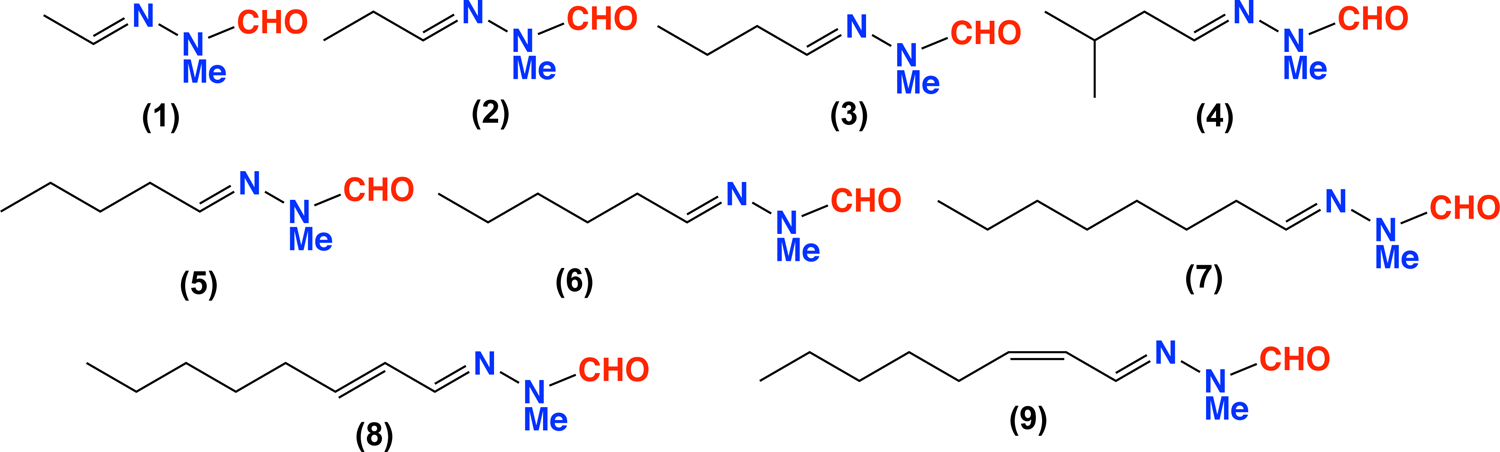
Chemical structures of gyromitrin (**1**) and its homologs (**2**–**9**).

**Figure 2.**
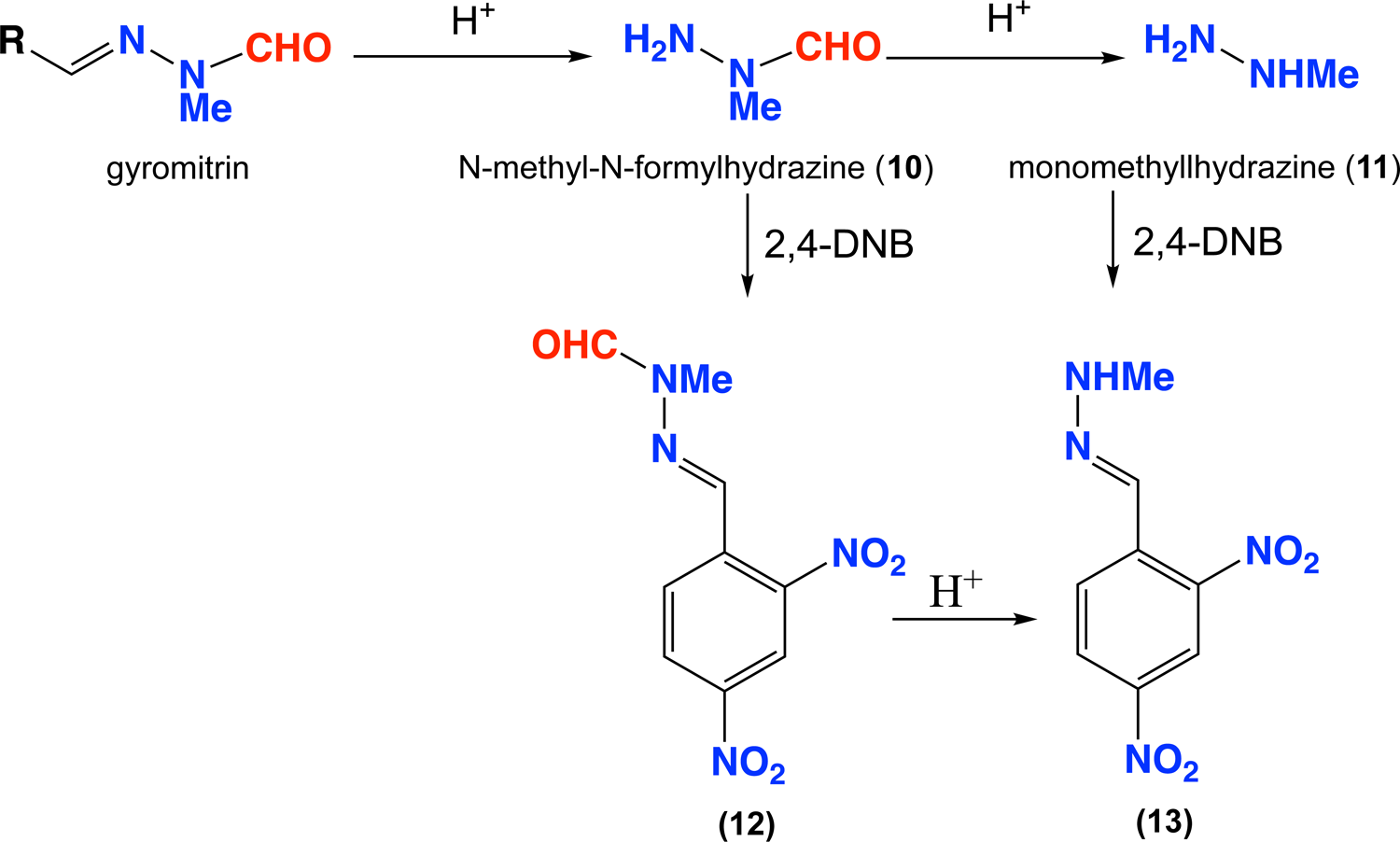
Acid hydrolysis of gyromitrin and homologs yields *N*-formyl-*N*-methylhydrazine (**10**) and monomethylhydrazine (**11**). Derivatization of **10** and **11** with 2,4-dinitrobenzaldehyde yields Schiff bases **12** and **13**.

Raw or undercooked *G. esculenta* mushrooms are poisonous primarily due to **10** and **11** formed via acid hydrolysis in the gastric environment. Symptoms usually appear from 5–12 hours after exposure. Most people poisoned by gyromitrin only experience gastrointestinal issues involving vomiting, abdominal pain, and diarrhea; in more severe cases, a victim suffers cytolytic hepatitis and jaundice (Mäkinen et al. 1977; Wright et al. 1978; Michelot and Toth 1991; Arłukowicz-Grabowska et al. 2019). Gyromitrin poisoning can also entail neurological symptoms such as vertigo, fatigue, and tremor, which can progress to seizure in the worst scenarios (Karlson-Stiber and Persson 2003). This is hypothesized to be a result of the binding of the gyromitrin hydrolytic products to pyridoxine (vitamin B_6_), inhibiting the enzymes that produce GABA and serotonin. Death, while rare, is caused by hepatic coma as unstable metabolic intermediates and methyl free radicals destroy the liver (Michelot and Toth 1991; Karlson-Stiber and Persson 2003; Arłukowicz-Grabowska et al. 2019). Gyromitrin may also have long-term toxic effects, including increased rates of cancer (Toth and Nagel 1978; Toth and Patil 1980). The genotoxic potential of gyromitrin informs the emerging hypothesis of its role in the development of neurodegenerative diseases such as sporadic amyotrophic lateral sclerosis (ALS) (Spencer 2020; Lagrange et al. 2021; Spencer and Kisby 2021; Spencer and Palmer 2021).

Beyond *Gyromitra esculenta*, whose toxicity has been well established, there is limited evidence regarding the toxicity of other *Gyromitra* species. At the turn of the 20^th^century, renowned mycophagist Charles McIlvaine regarded *G. brunnea*, *G. caroliniana*, and “*G. curtipes*” (*G. gigas* group) as “esculent” (McIlvaine and Macadam 1902). In 1976, *Gyromitra* taxonomist Harri Harmaja discussed a poisoning event in northern Sweden involving *G. ambigua* and concluded from the available evidence that this species contained dangerous quantities of gyromitrin in contrast to the nontoxic *G. infula* (Harmaja 1976). Harmaja also cited a report on four *Gyromitra* poisoning events in the Czech Republic, one of which supposedly involved *G. gigas*. However, this identification was based solely on its purported hefty stature and the specimen was never examined (Kubička 1966). A pattern emerges in older field guides whereby European *G. gigas* is flagged as toxic (references in Viernstein et al. [1980] and Weber [1995]) but species in the North American *G. gigas* group—now known to correspond to *G. korfii*, *G. montana*, and *G. americanigigas* (Miller et al. 2020, 2022)—are listed as edible (references in Weber [1995]). One such source, Tylutki’s Mushrooms of Idaho and the Pacific Northwest (1979), describes *G. montana* (listed as *G. gigas*) as a popular and delicious edible of the Rocky Mountains. From data gathered by the North American Mycological Association, *Gyromitra* spp. were the culprit behind about 4% of North American mushroom poisoning events involving humans between 1973 and 2005 (Beug et al. 2006; Beug 2014). *Gyromitra esculenta* was always involved when organ failure occurred. *Gyromitra brunnea* and *G. montana* resulted in poisonings as well, although symptoms appeared no more severe than those sometimes caused by other widely consumed mushrooms. In 2020, *G. venenata,* a species closely related to *G. esculenta*, was described from China after its consumption caused four people to go to the emergency room (Li et al. 2020). Finally, Lagrange et al. (2021) associated a hotspot of ALS in the French Alps to the consumption of lorchels, specifically *G. gigas*, although this species was perhaps prematurely implicated given that other unidentified *Gyromitra* species were also found stored in the residents’ homes.

Many different analytical chemistry techniques have been employed over the years to detect and/or quantify gyromitrin. After purifying and solving the chemical structure of gyromitrin, List and Luft (1968a, 1968b) developed spot tests involving acid hydrolysis of gyromitrin and subsequent derivatization of the hydrolytic products with various chromophores, as well as thin-layer chromatography (TLC) and potassium iodate titration methods. Pyysalo and colleagues developed gas chromatography (GC) methods to quantify gyromitrin hydrazone analogs or benzaldehyde derivatives of **10** and **11** (Pyysalo and Niskanen 1977; Pyysalo et al. 1978). Viernstein et al. (1980) also used GC along with an external standard calibration to quantify gyromitrin in the ether extracts of pressed mushrooms. Andary et al. (1984, 1985) separated ethanol extracts with TLC and sprayed *p*-dimethylaminocinnamaldehyde to form a red fluorophore quantifiable with spectrofluorometry. Finally, Arshadi et al. (2006) developed a GC-mass spectrometry analytical method involving the derivatization of acid-hydrolyzed ethanol extracts with pentafluorobenzoyl chloride. Despite all these methods, lacking from the literature is a method that employs high-performance liquid chromatography (HPLC) or ultra high-performance liquid chromatography (UHPLC) coupled with diode array detector (DAD), which are popular chromatographic techniques in modern chemistry labs.

Given this variety of published analytical methods, data on the presence of gyromitrin in lorchels is surprisingly scarce. In the first gyromitrin analysis outside of *G. esculenta*, Viernstein et al. (1980) detected small quantities of gyromitrin in two out of three European *G. gigas* specimens (0.05 and 0.74 mg/kg fresh specimen) but not in the single *G. fastigiata* specimen tested. Andary et al. (1985) did not detect gyromitrin in *Gyromitra perlata* or in Morchellaceae taxa but claimed to detect it in a curious set of species belonging elsewhere in the Pezizomycetes as well as the Leotiomycetes. Chemists at a toxicology laboratory in Michigan reported in a clinical toxicology conference abstract that *G. caroliniana* contained “minute” amounts of free **11** (Liang et al. 1998). On the other hand, in her popular morel hunting guide, Weber (1995) cited unpublished spot test data from Dr. Kenneth Cochran indicating the absence of gyromitrin in *G. caroliniana* as well as *G. brunnea* and *G. korfii*.

With the growing popularity of lorchel consumption (as evidenced by online forums such as the Facebook group *False Morels Demystified*), there is an urgent need to refine our understanding of which taxa contain the gyromitrin mycotoxin. Taking into consideration the available toxicity evidence, we hypothesized that gyromitrin evolved in the last common ancestor of *G. esculenta* and *G. ambigua*, with a loss of function in the closely related *Hydnotrya* clade due to the negative fitness consequences that mycotoxin production could have on hypogeous fungi relying on mycophagy for spore dispersal. To evaluate the distribution of gyromitrin in Discinaceae, we developed a simple, sensitive analytical method involving in situ acid hydrolysis of gyromitrin and chemical derivatization of its hydrolytic products, **10** and **11**, using 2,4-dinitrobenzaldehyde into Schiff bases **12** and **13** (FIG. 2). This derivatizing agent was chosen due to its proven efficiency in detection of related hydrazine-containing metabolites (Mohamed et al. 2018, 2021). Since the amount of gyromitrin is known to differ within the same species by environment and genotype (Marjatta and Pyysalo 1978; Andary et al. 1985), we aimed for broad taxon sampling and qualitative evaluation of gyromitrin presence rather than attempting to make claims about gyromitrin levels among different specimens within a species.

## MATERIALS AND METHODS

### Specimen acquisition.—

Ascocarp specimens were collected fresh by ACD or donated by mycologists and community scientists from across North America. Some collections were split with one half preserved at – 80 C and the other half dried in an electric dehydrator. All freshly collected dried specimen vouchers were deposited at the University of Michigan fungarium, except for a few that were entirely consumed through the gyromitrin detection method or did not dry properly. A subset of specimens was successfully cultured by sampling a piece of hymenium, placing it in sterile water, agitating the tissue to release ascospores, plating serial dilutions of the ascospore liquid on potato dextrose agar (PDA) and malt extract agar (MEA) with penicillin and streptomycin, and isolating single germinating ascospores onto PDA and MEA without antibiotics. In addition, a set of *Gyromitra* cultures from a recent study on *Pezizales* fungi were provided by Dr. A. Elizabeth Arnold from the living collection at the Robert L. Gilbertson Mycological Herbarium at the University of Arizona (Healy et al. 2022). Two *Gyromitra* cultures whose genomes have been sequenced were also purchased from the Westerdijk Fungal Biodiversity Institute CBS culture collection. Older dried ascocarp specimen vouchers were acquired as loans from the University of Michigan (MICH), University of Florida (FLAS), and University of Arizona (ARIZ) fungaria. Specimen vouchers and cultures tested for gyromitrin for this study are listed in TABLE 1.

**Table 1.**
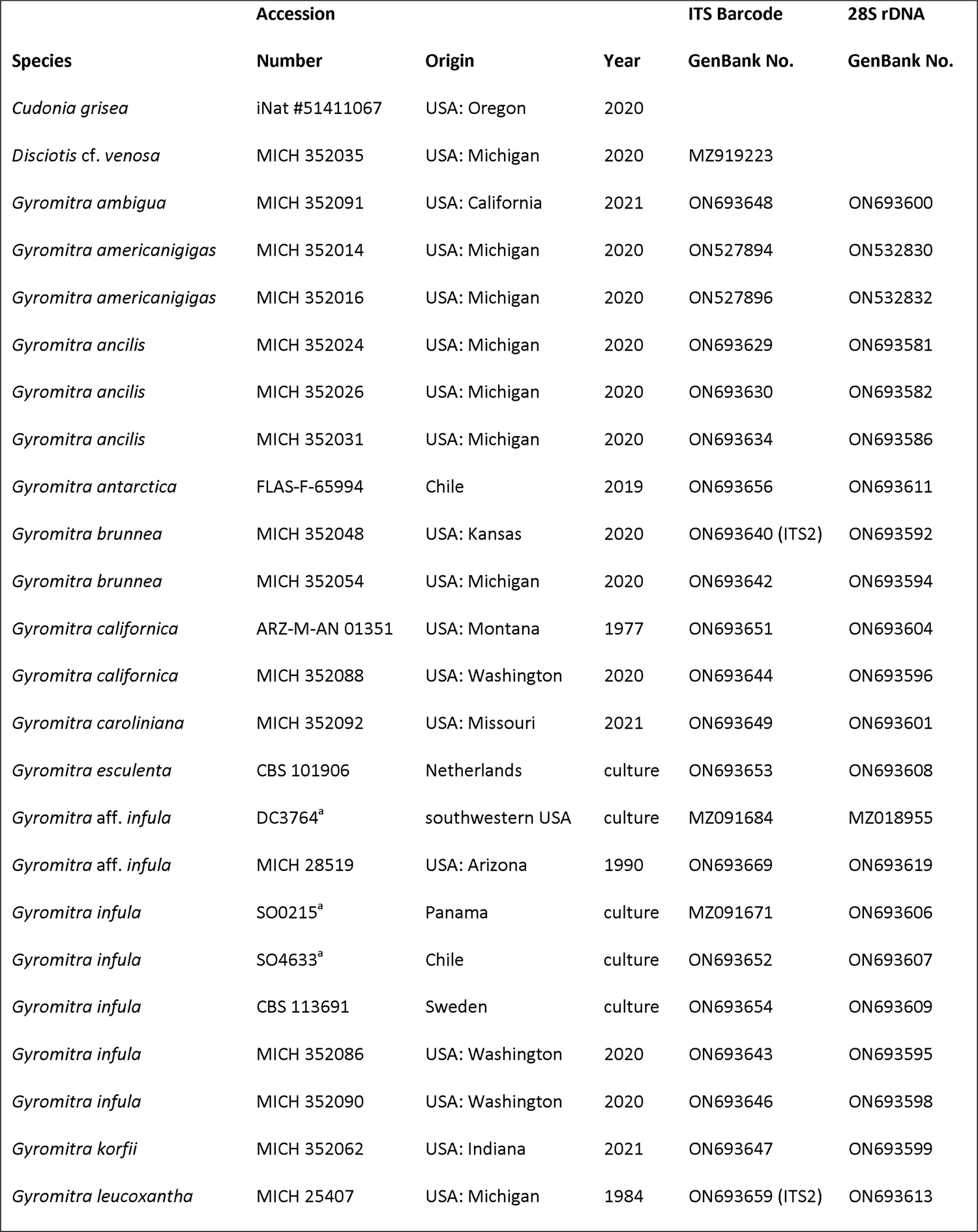
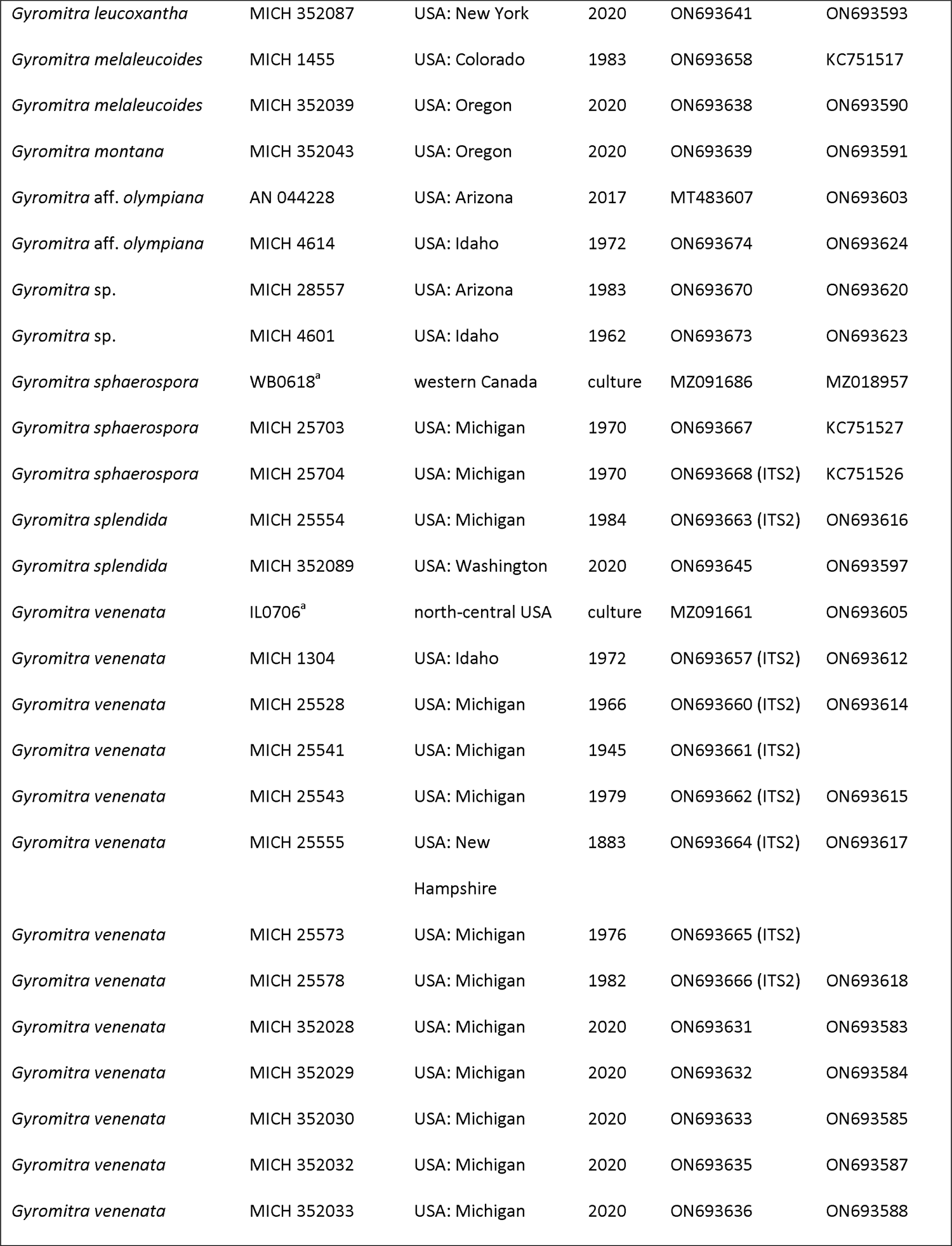
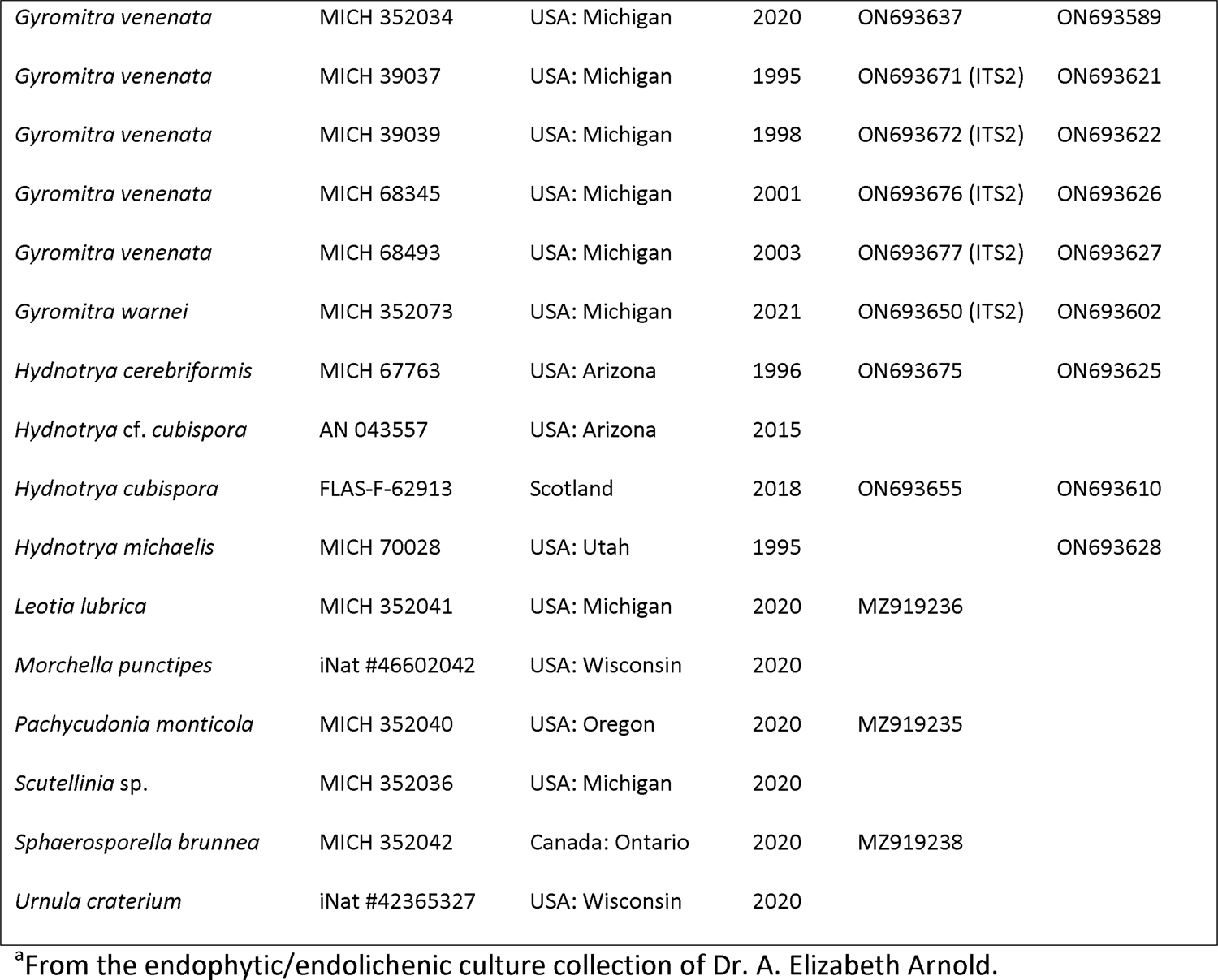
Specimens tested for gyromitrin and their associated metadata.

### Molecular data.—

DNA was extracted from dried ascocarps and fresh cultures using the 2 × CTAB method (Porter et al. 2011). Ascocarp tissue (approximately 0.25 cm^2^piece of hymenium) or hyphae (approximately 1 cm^2^scraped from mycelium growing on cellophane) were placed into a 1.5 mL microcentrifuge tube with 500 µL cell lysis solution (2 × CTAB extraction buffer: 2% cetyltrimethylammonium bromide, 1.4 M NaCl, 50 mM Tris, 10 mM Na_2_EDTA pH 8). Each sample was ground with a Kontes plastic pestle (DWK Life Sciences, England) for up to 2 min or until the tissue appeared homogenized, vortexed, and incubated in a water bath at 65 C for 60 min. Extraction proceeded with the addition of an equal volume of chloroform-isoamyl alcohol (24:1), centrifugation at 13 000 RPM (15 871 RCF) for 12 min, and transfer of the upper aqueous layer to a new 1.5 mL microcentrifuge tube. This extraction procedure was repeated once more and the final aqueous phase was precipitated with two-thirds volume ice cold isopropyl alcohol, mixed by inversion, and placed in a –20 C freezer overnight. The following day, the DNA was pelleted by centrifugation for 7 min. The alcohol supernatant was discarded, and the pellet rinsed with 1 mL cold 70% ethanol. The DNA was dried in a laminar flow hood and resuspended in 50 µL distilled water.

For each extraction, we attempted to amplify separately the nuc rDNA ITS1-5.8S-ITS2 (ITS barcode) region using the primers ITS1f and ITS4 (White et al. 1990; Gardes and Bruns 1993) and the D1–D2 domains of nuc 28S rDNA with the primers LR0R and LR5 (White et al. 1990; Moncalvo et al. 2000). If ITS amplification failed, we attempted to amplify the nuc rDNA ITS2 region using the primers ITS3 and ITS4 (White et al. 1990). PCR amplifications were carried out using the GoTaq Green Master Mix (Promega, Wisconsin) in a reaction volume of 12.5 µL with 5 µL of a 1:20 dilution of the DNA extract as a template. Reactions were completed on an Eppendorf Mastercycler Pro S Model 6325 thermal cycler. For ITS and ITS2, the thermal cycler parameters were as follows: 2 min initial denaturation at 94 C; 35 cycles of 30 s at 94 C, 30 s at 55 C, and 30 s at 72 C; and a 10 min final extension at 72 C. For 28S, they were: 5 min initial denaturation at 95 C; 40 cycles of 30 s at 95 C, 15 s at 52 C, and 1 min at 72 C; and a 10 min final extension at 72 C. Successful PCRs were enzymatically cleaned with ExoSAP-IT (Applied Biosystems, Massachusetts) and submitted to Genewiz for Sanger Sequencing (Azenta Life Sciences, New Jersey) using the PCR primers. Forward and reverse Sanger sequences were assembled with Geneious Prime (Dotmatics, Massachusetts) and submitted to GenBank (ON693581–ON693677) (TABLE 1).

### Phylogenetic analyses.—

Published Discinaceae ITS and LSU sequences were downloaded from GenBank with ENTREZ DIRECT (Kans 2020) (TABLE S1). Discinaceae ITS and LSU sequences were aligned separately with MUSCLE as implemented in SEAVIEW 5.0.4 (Edgar 2004; Gouy et al. 2010). Ambiguous regions of the alignments were trimmed with GBLOCKS 0.91b using less stringent parameters (Castresana 2000; Talavera and Castresana 2007) and concatenated with CATFASTA2PHYML 1.1.0 (github.com/nylander/catfasta2phyml). A maximum likelihood (ML) analysis was conducted on the concatenated alignment with RAXML 8.2.11 using the GTR+Gamma+I model of substitute evolution and 1000 bootstrap replicates (Stamatakis 2014; Abadi et al. 2019). Clades with bootstrap values ≥ 70% were considered significant and strongly supported (Hillis and Bull 1993). Bayesian analyses were performed with the same alignment under the above model using MRBAYES 3.2.7 on the CIPRES Science Gateway 3.3 portal (Miller et al. 2011; Ronquist et al. 2012). The Bayesian analyses lasted until the average standard deviation of split frequencies was below 0.01 with trees saved every 1000 generations and burn-in set at 25%. Bayesian posterior probabilities (BPP) were determined from a consensus tree using Geneious Prime. Clades with BPP ≥ 95% were considered significant and strongly supported (Larget and Simon 1999; Alfaro et al. 2003). The best ML tree was visualized and rooted with *Morchella esculenta* (Morchellaceae) as the outgroup in FIGTREE 1.4.4 (github.com/rambaut/figtree). The alignments and phylogenetic tree were deposited in Figshare (to be published upon manuscript acceptance).

### Gyromitrin analytical method development.— General experimental details

2,4-dinitrobenzaldehyde (2,4-DNB) and trifluoroacetic acid (TFA) were purchased from Sigma-Aldrich (D193607 and 302031). Authentic gyromitrin standard was purchased from Toronto Research Chemicals (G931900). HPLC-grade solvents were used in the gyromitrin extractions. Solvents used for HPLC, UHPLC, and HPLC-MS purposes were of HPLC-grade supplied by Labscan or Sigma-Aldrich and filtered/degassed through a 0.45 μm polytetrafluoroethylene membrane prior to use. Preparative HPLC was performed using Shimadzu LC-20AT HPLC instruments with corresponding detectors, fraction collectors, and software. Electrospray ionization mass spectra (ESIMS) were acquired using the Shimadzu LC-20AD Separations Module equipped with a Shimadzu LCMS-2020 Series mass detector in both positive and negative ion modes under the following conditions: Phenomenex Kinetex C8 1.7 µm 100 Å column, 50 µm × 2.1 mm, eluting with 0.4 mL/min of isocratic 90% H_2_O/acetonitrile (MeCN) for 1 min followed by gradient elution to 100% MeCN (with isocratic 0.1% formic acid modifier) over 6 min, at 210, 254, 280, and 370 nm. UHPLC-quadrupole time-of-flight (UHPLC-QTOF) analysis was performed on an UHPLC-QTOF instrument comprising an Agilent 1290 Infinity II UHPLC under the following conditions: Phenomenex Kinetex 1.7 μm phenylhexyl column, 50 µm × 2.1 mm, eluting with 0.4 mL/min of isocratic 90% H_2_O/MeCN for 1 min followed by gradient elution to 100% MeCN over 6 min (with isocratic 0.1% formic acid modifier) coupled to an Agilent 6545 LC/Q-TOF-MS system operating in positive mode, monitoring a mass range of 100 to 2000 amu.

### Synthesis of 2,4-dinitrobenzaldehyde Schiff base (13) reference compound

An aliquot of 2,4-DNB (94 mg, 0.48 mmol) dissolved in 4 mL ethanol was treated with **11** (120 mg, 2.67 mmol) and stirred at room temperature for 2 h. Afterwards, the reaction mixture was quenched with water (20 mL), extracted with 15 mL dichloromethane (CH_2_Cl_2_) two times, and the combined organic layer was dried over anhydrous sodium sulfate (Na_2_SO_4_), concentrated in vacuo, and purified by preparative reversed phase HPLC (Phenomenex Luna phenylhexyl, 21.2 mm × 25 cm, 5 μm, 20 mL/min, isocratic elution with 90% H_2_O/MeCN for 2 min followed by gradient elution from 90% H_2_O/MeCN to 100% MeCN over 30 min then isocratic elution with 100% MeCN for 5 min) to yield pure **13** (9 mg). Purity was checked by UHPLC coupled with diode array detection and electrospray ionization mass spectrometry (UHPLC-DAD-ESIMS) (FIG. S1). A seven-point calibration curve with solutions of reference compound **13** (0.39–50 µg/mL) in 50% H_2_O/MeCN was established by UHPLC-DAD analysis of an aliquot (10 µL) of each concentration in duplicate (FIG. S2). It is noteworthy that higher concentrations (100 µg/mL) of **13** did not fit into the calibration curve trendline due to saturation of the DAD detector.

### Detection of gyromitrin hydrazine hydrolytic products using 2,4-DNB

A series of gyromitrin standard solutions (5, 1, 0.5, 0.25, 0.1, 0.05, and 0.01 mg/mL) were prepared in 50% H_2_O/MeCN. An aliquot (10 μL) of each gyromitrin solution was transferred to glass vials containing 50% H_2_O/MeCN (400 μL) and treated with an aliquot (40 μL) of freshly prepared stock solution (5 mg/mL in MeCN) of 2,4-DNB and an aliquot (50 μL) of freshly prepared stock solution of 10% aqueous TFA. The reaction mixture was incubated at 40 C. Aliquots (10 μL) from each reaction mixture were analyzed by UHPLC-DAD at regular time intervals (0, 2, 5, 8, 13, 18, and 24 h) to detect gyromitrin hydrazine hydrolytic products derivatives with 2,4-DNB (FIG. S3).

### Detection of gyromitrin in Gyromitra venenata

Aliquots (50, 10 and 1 mg) of powdered *Gyromitra venenata* (MICH 352032) were transferred to glass vials containing 50% H_2_O/MeCN (820 μL) and treated with an aliquot (80 μL) of freshly prepared stock solution (5 mg/mL in MeCN) of 2,4-DNB and an aliquot (100 μL) of freshly prepared stock solution of 10% aqueous TFA. The reaction mixture was sonicated for 20 s and incubated at 40 C. Aliquots (10 μL) from each reaction mixture were analyzed by UHPLC-DAD at regular time intervals (0, 6, 13, and 24 h) to detect gyromitrin hydrazine hydrolytic products derivatives with 2,4-DNB. The reactions were performed in triplicate. Samples prepared by extracting the same amount of *G. venenata* with 50% H_2_O/MeCN (1000 μL) without the addition of 2,4-DNB and TFA were used as negative controls.

### Detection of gyromitrin in different samples

Gyromitrin analysis of samples proceeded as for *G. venenata* except for the following differences. Samples consisted of an aliquot of powdered ascocarp hymenium (ranging from 5– 100 mg, depending on available starting material) or a piece of mycelium scraped from a plate with cellophane (half a plate). An aliquot (not in triplicate) of the reaction mixture (10 µL) was analyzed after 13–18 h of incubation at 40 C. An equivalent amount of tissue was extracted with 50% H_2_O/MeCN (1000 μL) without the addition of 2,4-DNB and TFA as a negative control.

## RESULTS

UHPLC-DAD analysis of different concentrations of gyromitrin standard treated with 2,4-DNB and TFA with varying incubation times at 40 C revealed the need for 13–18 hours of incubation to complete the gyromitrin derivatization reaction to the monomethylhydrazine Schiff base (**13**). Our methodology was able to detect even trace amounts of gyromitrin to the extent of 10 ng in samples (FIG. S4). With a standard calibration curve of synthetic **13** (FIG. S2), the reaction recovery percentage was calculated based on peak areas of **13** obtained from derivatization of different gyromitrin standard concentrations (TABLE S2). The peak area of the monomethylhydrazine Schiff base (**13**) was plotted against the gyromitrin standard concentration showing a linear relationship (FIG. S5). This correlation could be used for the quantification of gyromitrin in fungal samples. Equipped with a sensitive gyromitrin analytical method, aliquots of dried *Gyromitra venenata* (MICH 352032) tissue were processed with different incubation times to determine the hydrolysis of gyromitrin present in ascocarp samples (FIG. 3). The peak area of Schiff base **13** remained relatively constant between 13 and 24 hours of extraction regardless of the amount of starting material (FIG. S6). A large amount of gyromitrin (2598 mg/kg dried ascocarp) was detected in these aliquots of *G. venenata*.

**Figure 3.**
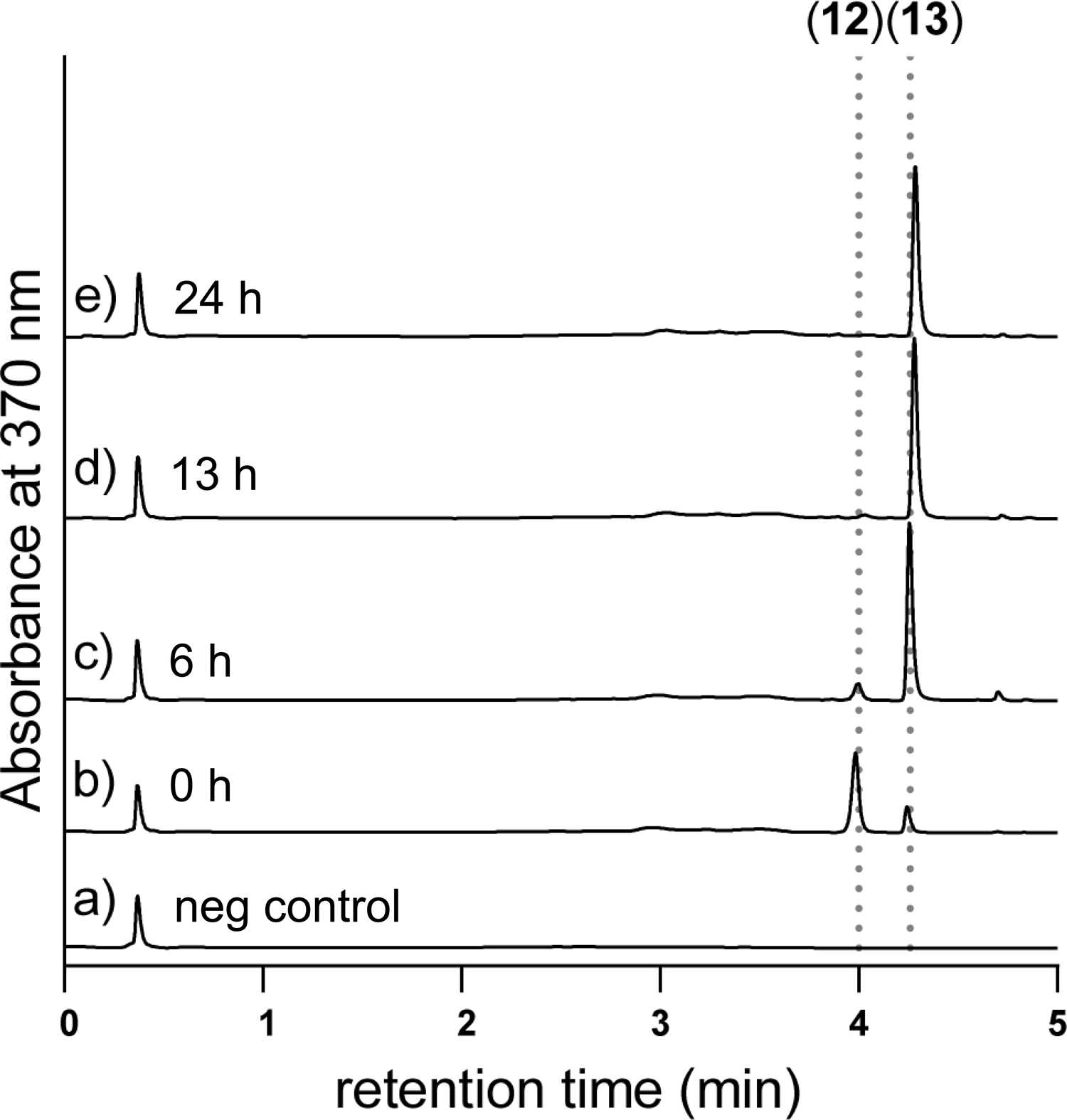
UHPLC-DAD (370 nm) chromatograms of derivatization progress of gyromitrin in 10 mg of dried *Gyromitra venenata* ascocarp (MICH 352032). A. Ascocarp extract without 2,4-DNB/TFA treatment (negative control). B–E. Ascocarp extract treated with 2,4-DNB/TFA with different incubation time intervals (0, 6, 13, and 24 h, respectively).

We analyzed 66 specimens as dried ascocarps, freshly frozen ascocarps, living cultures, or a combination thereof, resulting in 105 individual tests (TABLE S3). All taxa outside the lorchel family (eight specimens) tested negative for gyromitrin. The majority of Discinaceae taxa also tested negative for gyromitrin. Both *G. leucoxantha* specimens tested positive with MICH 352087 containing 55 mg/kg of gyromitrin per dried ascocarp (MICH 25407 was not quantified). The chromatograms also included a strong peak between **12** and **13**, which was absent from the negative control (extracts without the addition of 2,4-DNB and TFA), that likely corresponds to an unidentified Schiff base derivative produced by *G. leucoxantha* (FIG. 4). All four taxa in the *G. esculenta* group (*G. antarctica*, *G. esculenta*, *G. splendida,* and *G. venenata*) tested positive for gyromitrin, with a few caveats. First, gyromitrin levels generally decreased with age of the dried ascocarp, which could be attributed to the volatile property of gyromitrin. Still, even the oldest voucher tested (*G. venenata* [MICH 25555] from 1883) had trace levels of gyromitrin in the chromatogram (FIG. 5). Second, cultures growing on PDA tested positive, but gyromitrin was not detected in cultures growing on MEA (TABLE S3). Third, the two *G. esculenta* group ascocarps from the western United States contained relatively small amounts of gyromitrin. *Gyromitra splendida* (MICH 352089) from Washington state contained 92 mg/kg of gyromitrin per dried ascocarp (FIG. 4). *G. venenata* (MICH 1304) from Idaho also appeared to have less gyromitrin than expected, although this is based solely on the relative height of peaks in the chromatogram (FIG. 5d). Evidence for the toxicity and gyromitrin content of *Gyromitra* species is summarized in TABLE 2.

**Figure 4.**
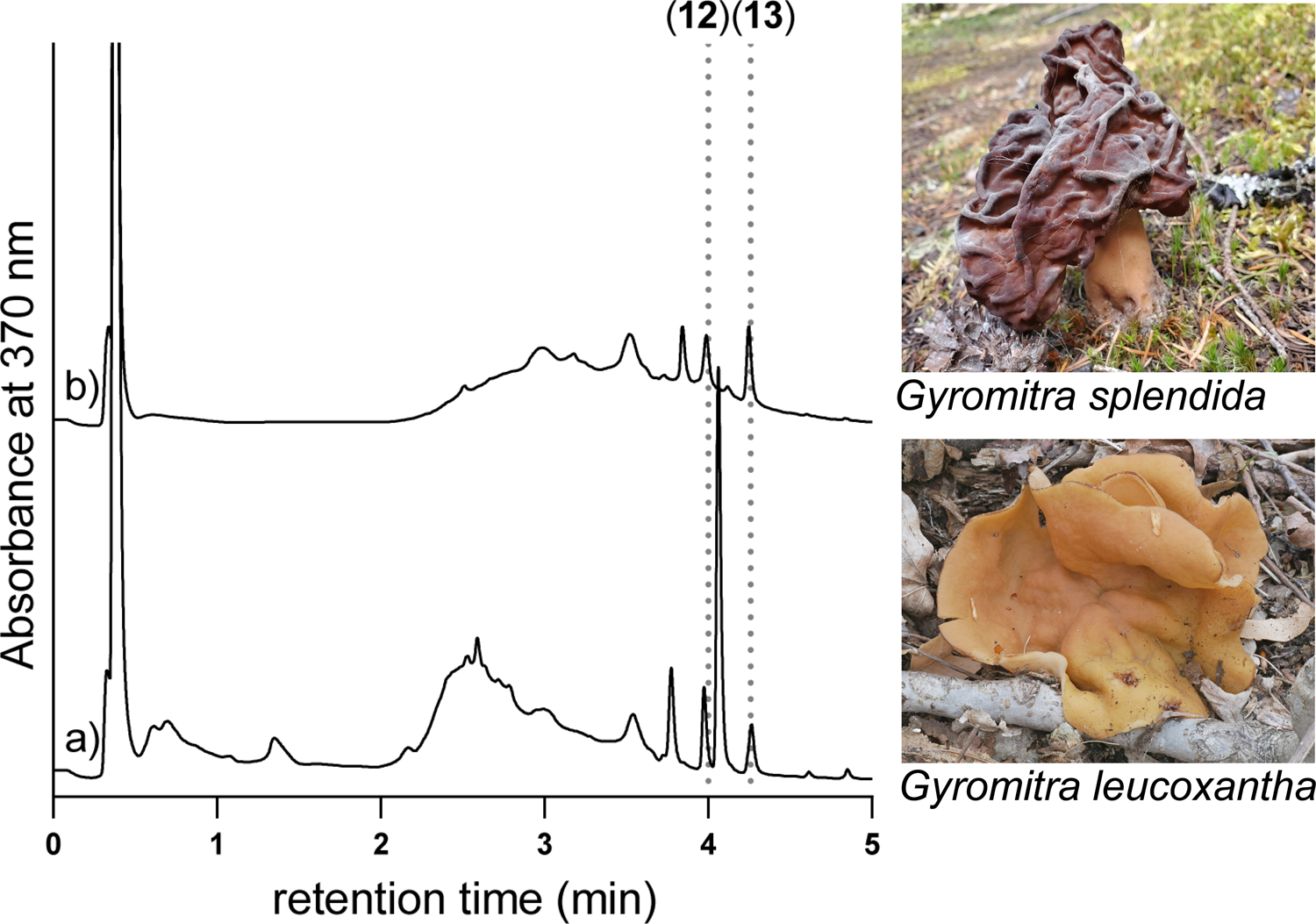
UHPLC-DAD (370 nm) chromatograms of ascocarp extracts. A. *Gyromitra leucoxantha* (MICH 352087). B. *Gyromitra splendida* (MICH 352089). Photograph of *G. splendida* by Eric and Jen Chandler. Photograph of *G. leucoxantha* by Garrett Taylor.

**Figure 5.**
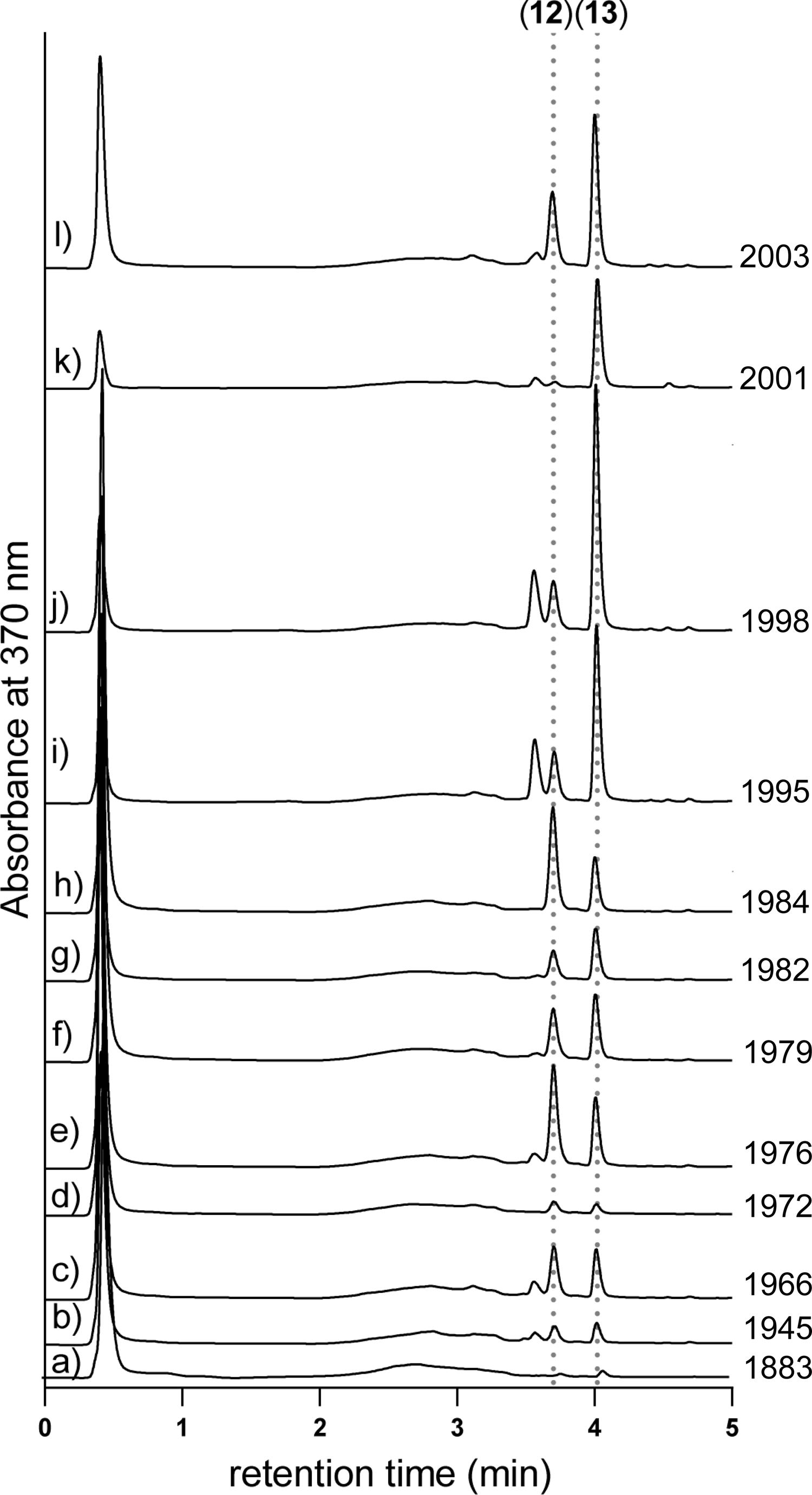
UHPLC-DAD (370 nm) chromatograms of 100 mg of *Gyromitra venenata* (A–G, I–L) or *Gyromitra splendida* (H) dried ascocarps collected in different years. A. MICH 25555 (1883). B. MICH 25541 (1945). C. MICH 25528 (1966). D. MICH 1304 (1972). E. MICH 25573 (1976). F. MICH 25543 (1979). G. MICH 25578 (1982). H. MICH 25554 (1984). I. MICH 39037 (1995). J. MICH 39039 (1998). K. MICH 68345 (2001). L. MICH 68493 (2003).

**Table 2.**
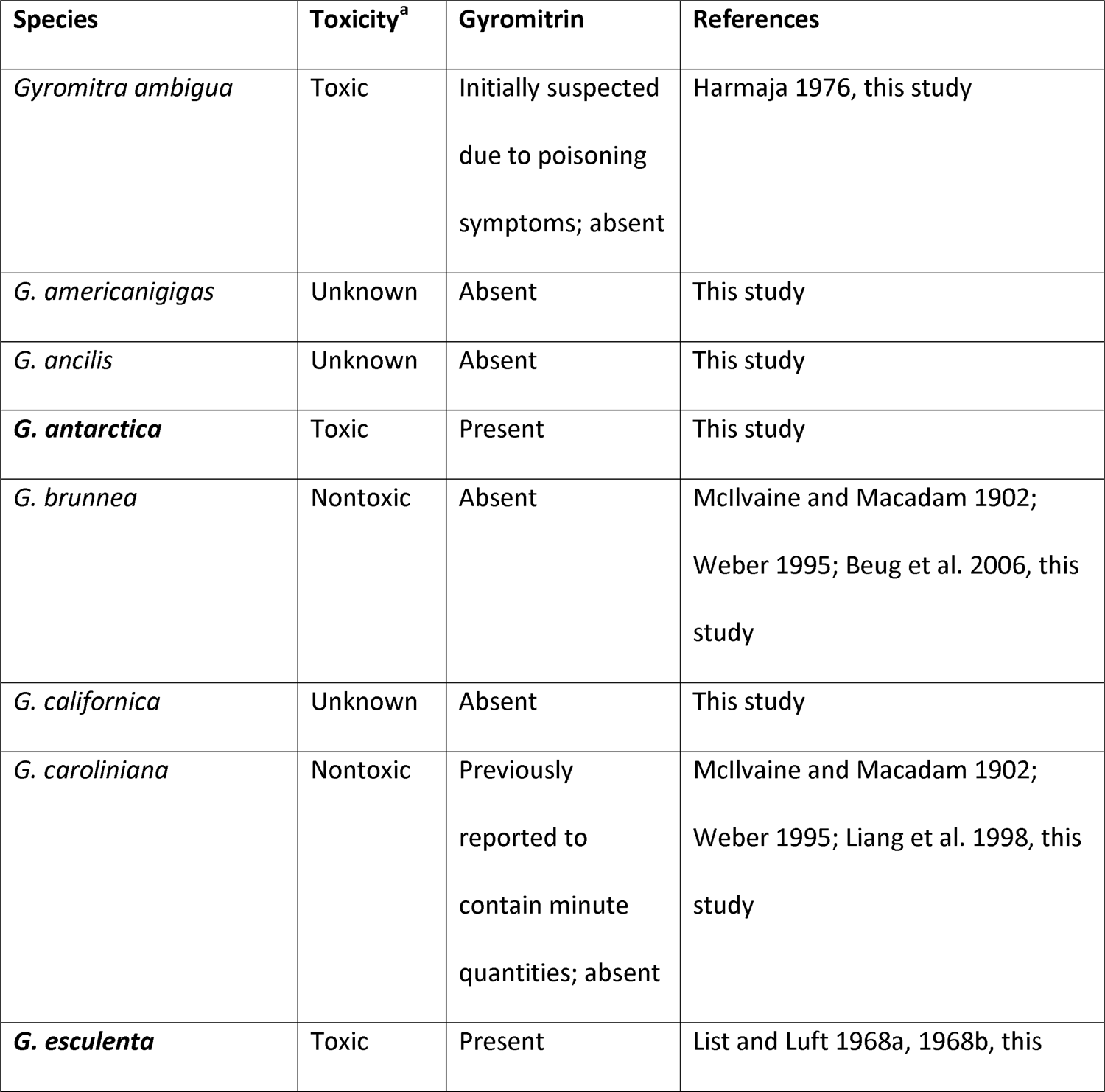
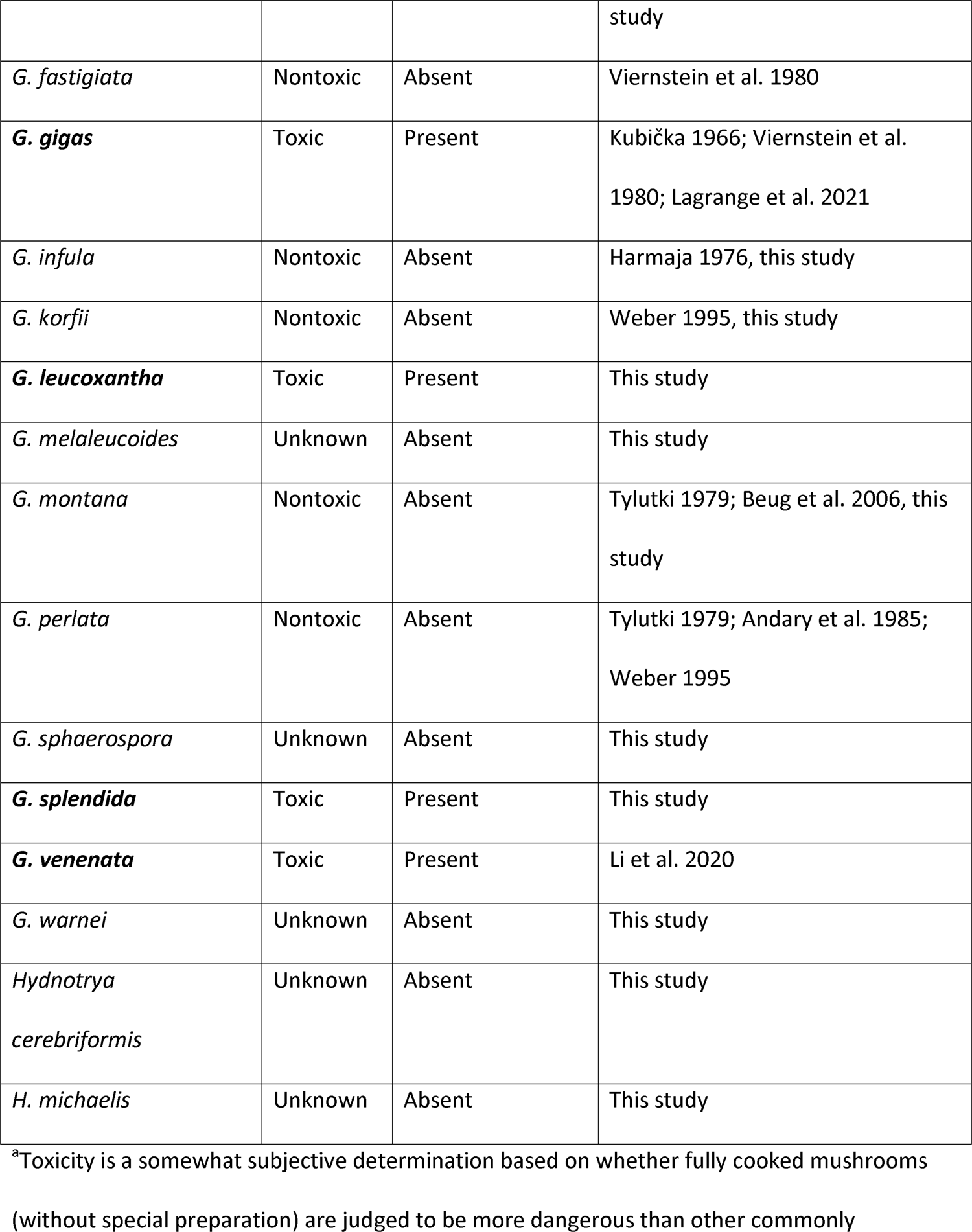
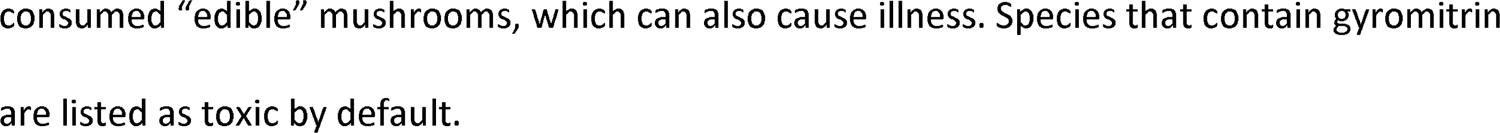
Summary of available evidence for the toxicity and presence of gyromitrin in Discinaceae species based on primary literature and tests conducted for this study. Bolded species are taxa that have been shown to contain gyromitrin. Evidence of toxicity does not necessitate that gyromitrin is the etiological factor, and no evidence of toxicity does not mean gyromitrin is absent from that species.

We had high success in sequencing rDNA from Discinaceae specimens, including from old fungarium vouchers—56/58 specimens (97%) had either the entire ITS or ITS2 region successfully sequenced and 54/58 specimens (93%) had the 28S rDNA barcode successfully sequenced. *Hydnotrya* specimens proved to be the most challenging to sequence, perhaps because of the contamination associated with their subterranean habitat. The concatenated ITS and 28S rDNA alignment contained 86 sequences with 1402 characters and 571 distinct patterns, 387 of which were parsimony informative. The maximum likelihood phylogenetic tree for the concatenated dataset allowed us to validate species determinations and visualize gyromitrin content in a phylogenetic context (FIG. 6). Across Discinaceae, a general absence of gyromitrin was punctuated by its discontinuous presence in two separate groups, the *G. esculenta* and *G. leucoxantha* clades. Due to the use of just two loci, bootstrap support for basal nodes was generally low and the evolutionary relationships between well-defined clades remains ambiguous.

**Figure 6.**
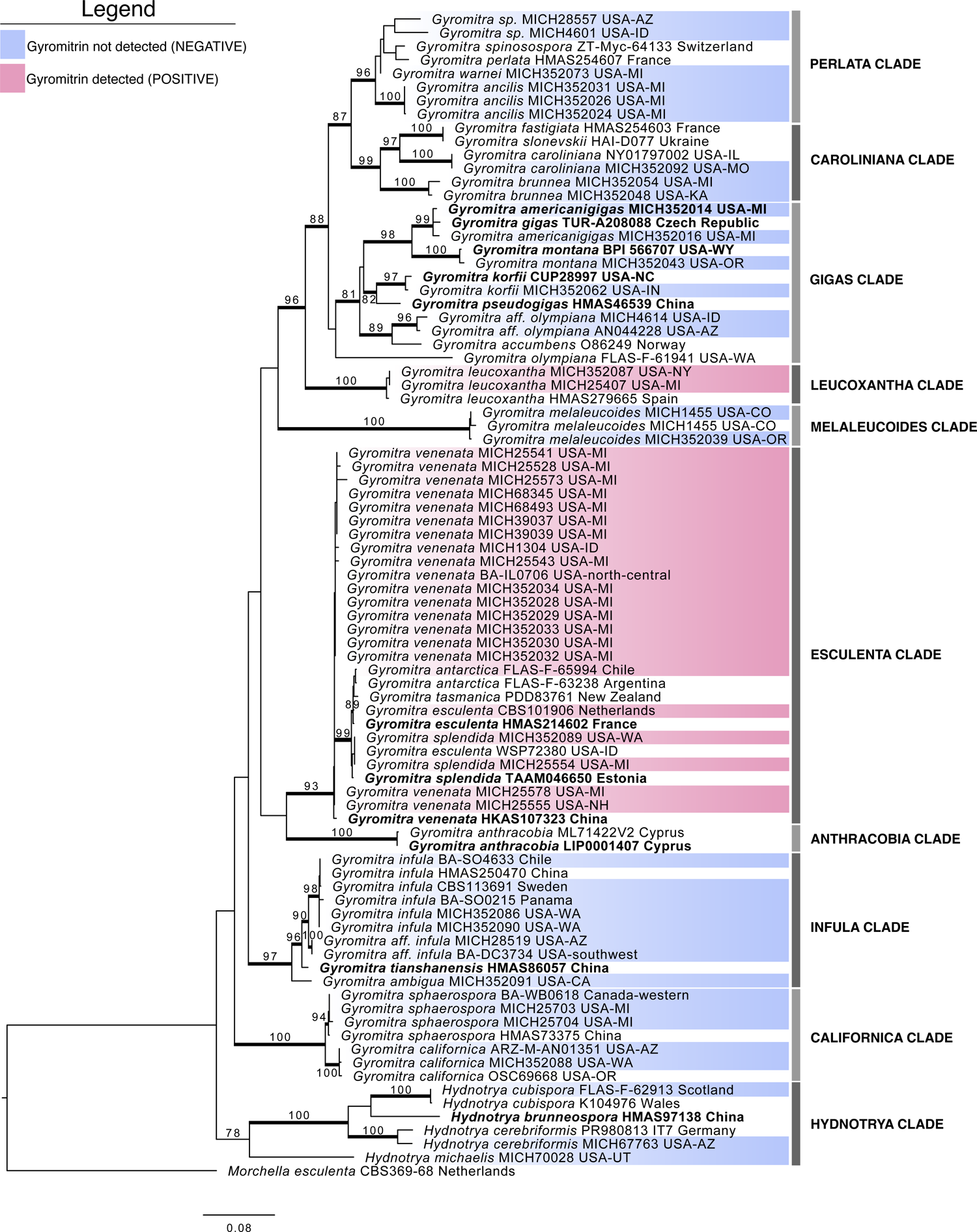
Discinaceae maximum likelihood phylogenetic tree based on concatenated aligned sequences of the ITS barcode and 28S rDNA showing the presence and absence of gyromitrin. Specimens tested for gyromitrin are highlighted: blue means no gyromitrin was detected (negative); red means gyromitrin was detected (positive). Bootstrap values ≥ 0.70 are shown at the nodes and Bayesian posterior probability scores above 0.95 are indicated by thickened branches. Taxon names for sequences derived from taxonomic type material are bolded.

## DISCUSSION

Gyromitrin determination using our newly developed UHPLC analytical method indicated that gyromitrin production is concentrated in the *G. esculenta* group, represented here by *G. antarctica*, *G. esculenta*, *G. splendida*, and *G. venenata* (although taxonomic names will likely change when the group is treated in a modern systematic revision). Contrary to expectations, we failed to detect gyromitrin in *G. ambigua* or *G. infula,* but to our surprise we detected gyromitrin in *G. leucoxantha*. Accepting at face value earlier reports of gyromitrin production by *G. gigas* sensu stricto (Viernstein et al. 1980), six loss events are required to explain this pattern according to our phylogeny (*G. melaleucoides* clade, last common ancestor of *G. caroliniana* and *G. ancilis* clades, and various losses in the *G. gigas* clade). A similar number of loss events, or one horizontal transfer and four loss events, would be required to explain this distribution according to previously published phylogenetic trees (Wang and Zhuang 2019; Miller et al. 2020, 2022). On the other hand, only two horizontal transfer events could result in the observed distribution, making this the most parsimonious hypothesis. However, repeated occurrences of genetic loss of function have been documented elsewhere (Patron et al. 2007; Morris et al. 2012), so we are unable to infer a definitive history of the evolution of gyromitrin in Discinaceae.

The distribution of gyromitrin observed here is consistent with a model of rapid evolution coupled with horizontal transfer, which is typical for secondary metabolites like gyromitrin (Rokas et al. 2020). In fungi, secondary metabolites—small, bioactive molecules not required for growth but important for interspecific interactions—are usually produced by enzymes encoded by genes physically clustered in the genome as a biosynthesis gene cluster (BGC). BGCs can experience strong selective pressure due to their importance in ecological interactions that often results in a narrow taxonomic distribution as well as intraspecific variation in the ability to produce a given secondary metabolite (Rokas et al. 2020; Yancey et al. 2022). Horizontal gene transfer is also a powerful evolutionary force that has led to the broad and phylogenetically disjunct expression of mycotoxins such as amanitin, psilocybin, and epipolythiodioxopiperazines (Patron et al. 2007; Rokas et al. 2020; Luo et al. 2022; Van Court et al. 2022). Therefore, a disjointed distribution of gyromitrin is not unprecedented.

The UHPLC-DAD analytical method presented in this manuscript is an accessible method that could facilitate further investigations into the gyromitrin mycotoxin, including determinations in human body fluids. While other methods like spot tests and TLC are simpler, they have low detection sensitivity, accuracy, and resolution. Thus, earlier reported conclusions of Andary et al. (1985) that taxonomically disparate ascomycete fungi contain gyromitrin should be reinterpreted. These taxa likely contain hydrazines or natural products that hydrolyze to form hydrazines, but not necessarily gyromitrin. Hydrazines belong to a class of chemicals defined by their nitrogen-nitrogen bond (N-N bond), of which only about 200 have been characterized to date, or less than 0.1% of the total known natural products (Blair and Sperry 2013; Le Goff and Ouazzani 2014). The frequency at which hydrazine derivatives were detected by Andary et al. (1985) suggest they might be more prevalent in fungi than expected and indicate that Pezizomycetes and Leotiomycetes fungi should be broadly screened for secondary metabolites containing an N-N bond.

### On the edibility of lorchels.—

Given that many people are interested in consuming lorchels and that our research will undoubtedly influence the conversation around their edibility, we believe it is pertinent that we weigh in on the matter. While the methods exist to eat gyromitrin-containing lorchels without acute poisoning, the dangers of improperly preparing them and their potential link to neurodegenerative disease lead us to conclude that the *Gyromitra esculenta* group should never be consumed. The same is true for *G. leucoxantha*, which we demonstrated to have the capacity to produce gyromitrin, albeit in relatively small quantities. With the gross morphology of a cup fungus, there is no culture of eating this species and it would be best for it to stay that way (Weber 1995). As a potential producer of gyromitrin, we recommend individuals avoid European *G. gigas* as well, at least until its status is clarified with contemporary analysis (Viernstein et al. 1980). We acknowledge that people are likely to continue ingesting some of these species, in which case it is crucial to cook them according to the safety guidelines established for their commercial sale in Europe, like that of the Finnish Food Safety Authority (Sitta et al. 2021).

Outside of these taxa, we believe one’s decision to consume lorchels is a matter of personal preference based on an informed consideration of the risks involved. In favor of some North American lorchels, there is already an established and widespread culture of consuming *G. brunnea*, *G. caroliniana*, *G. korfii*, *G. montana*, and presumably *G. americanigigas*, and no evidence of extraordinary toxicity or gyromitrin production—except for a curious, unpublished report of “minute” quantities of free **11** in *G. caroliniana* that our data did not corroborate (Weber 1995; Liang et al. 1998; Beug et al. 2006; Beug 2014). With these lorchels one should adhere to the central tenants of all wild mushroom foraging: positively identify the species, only consume fresh mushrooms (not mushrooms with signs of insect damage, moldy growth, or putrescence), thoroughly cook the mushrooms, try a little before trying a whole meal to evaluate idiosyncratic responses like an allergic reaction, and be careful with overindulgence.

In opposition to their consumption, we likely have not established the full distribution pattern of gyromitrin in Discinaceae. Intraspecific variation in the amount of gyromitrin produced by *G. gigas* resulted in one specimen having no detectable levels (Viernstein et al. 1980), meaning even some species that tested negative in this study might have the capacity to produce gyromitrin. Furthermore, *Gyromitra* species have up to a dozen biosynthesis gene clusters whose corresponding secondary metabolites are entirely uncharacterized (MycoCosm: Grigoriev et al. 2014). Given that each species could produce a unique set of secondary metabolites, there are likely hundreds of undescribed chemicals with potentially powerful bioactive properties and implications for human health across the lorchel family. For example, the characterization of 1-(2-hydroxyacetyl)-pyrazol in *G. fastigiata*, a chemical with structural similarity to gyromitrin, might give pause to consumers of *G. fastigiata* as well as its sister species *G. caroliniana* (Jurenitsch et al. 1988). In rebuttal, one could point out that this is the case for almost every mushroom that is regarded as edible. For example, choice edible morels (*Morchella* spp.) can occasionally cause illness, including gastrointestinal distress, vomiting, diarrhea, and sometimes even neurological effects, yet the nature of the toxin is a complete mystery (Benjamin 2015). Even when a species contains a potentially toxic substance, like agaritine in the common button mushroom (*Agaricus bisporus*), it may still be consumed because the quantities present are generally recognized as safe *enough* for humans (Roupas et al. 2010; Khovpachev et al. 2021). Regardless, due to the uncharacterized chemodiversity in Discinaceae, unchartered species should be treated with extreme caution, even when a species is closely related to others that are consumed without issue.

### Conclusions and open questions.—

Our study is the most extensive sampling to date of lorchels for their capacity to produce gyromitrin. Further sampling will only improve our resolution of the evolution of gyromitrin production in Discinaceae. Along with *G. gigas*, it remains an open question whether *G. anthracobia* produces gyromitrin. Described in 2018 from Cyprus, *G. anthracobia* has the closest phylogenetic affinity to the *Gyromitra esculenta* group and thus is considered suspect (Crous et al. 2018; Li et al. 2020). Beyond gyromitrin, explorations of lorchel chemodiversity would not only reveal novel and potentially useful natural products but could also shed light on the etiology of poisonings caused by gyromitrin-free taxa, like *G. ambigua*. The relationship, if any, between 1-(2-hydroxyacetyl)-pyrazol in *G. fastigiata* and gyromitrin is a critical question. The analytical method reported here has the potential to identify previously unreported hydrazine-containing toxins as shown in the strong, uncharacterized peak in the chromatograms of *G. leucoxantha* (FIG. 4). This finding deserves dedicated follow-up studies to structurally characterize the unknown entity.

The lower levels of gyromitrin produced by western *G. esculenta* group ascocarps are also an avenue for future study. Tylutki (1979) mentions that *“G. esculenta*” has been eaten by many people in the western United States without ill effect, hypothesizing that mushrooms in this region contain less gyromitrin than those in the east (although poisonings are not unheard of [Leathem and Dorran 2007]). *Gyromitra splendida* (MICH 352089) from Washington had low levels of gyromitrin (28 × less than *G. venenata* [MICH 352032] from Michigan), providing some support for this hypothesis. Gyromitrin levels seem to decrease with altitude (Andary et al. 1985), which could explain this pattern, but the full extent of the effect of gene-by-environment interactions on the gyromitrin phenotype is an open question. Future studies should systematically explore the differential production of gyromitrin on different media. Our finding that gyromitrin is produced on PDA but not MEA encourages the use of transcriptomics to identify genes whose expression is correlated with mycotoxin production, facilitating the identification of the gyromitrin biosynthesis gene cluster. Identification of the gyromitrin genes will ultimately allow for the elucidation of the evolution of gyromitrin biosynthesis, discovery of homologous gene clusters in Discinaceae and other fungal clades, and characterization of novel secondary metabolites with related chemical structures.

## ACKNOWLEDGEMENTS

We extend our gratitude to all the people who collected and donated specimens, which greatly augmented this project: Leah Bendlin, Eric Chandler, Jen Chandler, Jonathan Frank, Igor Khomenko, Kitty Lundeen-Ness, Bee Marcotte, Steve Ness, Dave Ramos, Jon Shaffer, Huafang Su, Garrett Taylor, Jud Vanwyk, and Jeff Volpert. We also thank the fungarium curators, collection managers, and assistants who supported this research by loaning specimen vouchers and cultures: Dr. A. Elizabeth Arnold, Joseph Myers, and Ming-Min Lee (ARIZ); Dr. Matthew Smith, Dr. Rosanne Healy, and Benjamin Lemmond (FLAS); and Patricia Rogers and Dr. Alison Harrington (MICH). The strains provided by Dr. A. Elizabeth Arnold were collected with support from the National Science Foundation under grants DEB-1045766 and DEB-1541496, with additional support from the University of Arizona’s College of Agriculture and Life Sciences and USDA NIFA awards ARZT-1361340-H25-242 and ARZT-1259370-S25-200. Michelle Orozco-Quime’s work in trialing gyromitrin spot tests is much appreciated. Vavřinec Klener provided a helpful translation of a section of Kubička (1966). Taylor M. Tai provided thoughtful editorial advice on earlier drafts of this manuscript. This research was supported by the Michigan Translational Research and Commercialization (MTRAC) Innovation Hub for AgBio and the Canadian Institute for Advanced Research (CIFAR). AT received funding from the University of Michigan Biological Science Initiative. TYJ is a fellow of the CIFAR research program “Fungal Kingdom: Threats & Opportunities”. The authors report there are no competing interests to declare.

## AUTHOR CONTRIBUTIONS

ACD and TYJ conceived of the study. ACD collected specimens and sequenced their DNA. ACD and ANM conducted the phylogenetic analyses. ANM provided molecular data to aid in species identification. OGM and AT designed and developed the gyromitrin analytical method. OGM carried out chemical synthesis and analyzed the UHPLC-DAD chemical profiles. ACD, OGM, and PS carried out gyromitrin analysis of mushroom samples. ACD and OGM drafted the manuscript and constructed the Supporting Information. All authors reviewed and agreed to the published version of the manuscript.

## Notes

### Competing Interest Statement

The authors have declared no competing interest.

## LITERATURE CITED

1. Abadi S, Azouri D, Pupko T, Mayrose I. 2019. Model selection may not be a mandatory step for phylogeny reconstruction. Nature Communications 10:1–11.

2. Alfaro ME, Zoller S, Lutzoni F. 2003. Bayes or bootstrap? A simulation study comparing the performance of Bayesian Markov chain Monte Carlo sampling and bootstrapping in assessing phylogenetic confidence. Molecular Biology and Evolution 20:255–266.

3. Andary C, Privat G, Bourrier M-J. 1984. Microdosage spectrofluorimétrique sur couches minces de la monométhylhydrazine chez Gyromitra esculenta. Journal of Chromatography 287:419– 424.

4. Andary C, Privat G, Bourrier M-J. 1985. Variations of monomethylhydrazine content in *Gyromitra esculenta*. Mycologia 77:259–264.

5. Arłukowicz-Grabowska M, Wójcicki M, Raszeja-Wyszomirska J, Szydłowska-Jakimiuk M, Piotuch B, Milkiewicz P. 2019. Acute liver injury, acute liver failure and acute on chronic liver failure: A clinical spectrum of poisoning due to *Gyromitra esculenta*. Annals of Hepatology 18:514–516.

6. Arshadi M, Nilsson C, Magnusson B. 2006. Gas chromatography-mass spectrometry determination of the pentafluorobenzoyl derivative of methylhydrazine in false morel (*Gyromitra esculenta*) as a monitor for the content of the toxin gyromitrin. Journal of Chromatography A 1125:229–233.

7. Benjamin DR. 2015. Neurological effects of *Morchella* sp. FUNGI 8:24–25.

8. Benjamin DR. 2020. Gyromitrin poisoning: More questions than answers. FUNGI 13:36–39.

9. Beug MW. 2014. False morels: Age-old questions of edibility. FUNGI 7:28–31.

10. Beug MW, Shaw M, Cochran KW. 2006. Thirty-plus years of mushroom poisoning: Summary of the approximately 2000 reports in the NAMA case registry. McIlvainea: Journal of American Amateur Mycology 16:47–68.

11. Blair LM, Sperry J. 2013. Natural products containing a nitrogen-nitrogen bond. Journal of Natural Products 76:794–812.

12. Castresana J. 2000. Selection of conserved blocks from multiple alignments for their use in phylogenetic analysis. Molecular Biology and Evolution 17:540–552.

13. Court RC Van, Wiseman MS, Meyer KW, Ballhorn DJ, Amses KR, Slot JC, Dentinger BTM, Garibay-Orijel R, Uehling JK. 2022. Diversity, biology, and history of psilocybin-containing fungi: Suggestions for research and technological development. Fungal Biology 126:308–319.

14. Crous PW, Wingfield MM, Burgess TI, Hardy GESJ, Gené J, Guarro J, Baseia IG, García D, Gusmão LFP, Thangavel R, Adamčík S, Barili A, Barnes CW, Bezerra JDP, Bordallo JJ, Cano-Lira JF, de Oliveira RJ V., Ercole E, Hubka V, Iturrieta-González I, Kubátová A, Martín MP, Moreau P-A, Morte A, Ordoñez ME, Rodríguez A, Stchigel AM, Vizzini A, Abdollahzadeh J, et al. 2018. Fungal Planet description sheets: 716–784. Persoonia 40:240–393.

15. Edgar RC. 2004. MUSCLE: Multiple sequence alignment with high accuracy and high throughput. Nucleic Acids Research 32:1792–1797.

16. Gardes M, Bruns TD. 1993. ITS primers with enhanced specificity for basidiomycetes— application to the identification of mycorrhizae and rusts. Molecular Ecology 2:113–118.

17. Goff G Le, Ouazzani J. 2014. Natural hydrazine-containing compounds: Biosynthesis, isolation, biological activities and synthesis. Bioorganic and Medicinal Chemistry 22:6529–6544.

18. Gouy M, Guindon S, Gascuel O. 2010. Sea view version 4: A multiplatform graphical user interface for sequence alignment and phylogenetic tree building. Molecular Biology and Evolution 27:221–224.

19. Grigoriev I V., Nikitin R, Haridas S, Kuo A, Ohm R, Otillar R, Riley R, Salamov A, Zhao X, Korzeniewski F, Smirnova T, Nordberg H, Dubchak I, Shabalov I. 2014. MycoCosm portal: Gearing up for 1000 fungal genomes. Nucleic Acids Research 42:D699–D704.

20. Härkönen M. 1998. Uses of mushrooms by Finns and Karelians. International Journal of Circumpolar Health 57:40–55.

21. Harmaja H. 1976. Another poisonous species discovered in the genus *Gyromitra*: *G. ambigua*. Karstenia 15:36–37.

22. Healy RA, Arnold AE, Bonito G, Huang YL, Lemmond B, Pfister DH, Smith ME. 2022. Endophytism and endolichenism in Pezizomycetes: The exception or the rule? New Phytologist 233:1974–1983.

23. Hillis DM, Bull JJ. 1993. An empirical test of bootstrapping as a method for assessing confidence in phylogenetic analysis. Systematic Biology 42:182–192.

24. Jurenitsch J, Aurada E, Kubelka W. 1988. 1-(2-Hydroxyacetyl)-pyrazol, eine neue, mit gyromitrin verwandte verbindung aus *Gyromitra fastigiata*. Zeitschrift Für Mykologie 54:155–157.

25. Kans J. 2020. Entrez Direct: E-utilities on the UNIX Command Line. In: Entrez Programming Utilities Help [Internet]. Bethesda, MD: National Center for Biotechnology Information. p. 1–65.

26. Karlson-Stiber C, Persson H. 2003. Cytotoxic fungi: An overview. Toxicon 42:339–349.

27. Khovpachev AA, Basharin VA, Chepur S V., Volobuev S V., Yudin MA, Gogolevsky AS, Nikiforov AS, Kalinina LB, Tyunin MA. 2021. Actual concepts of higher fungi’s toxins: Simple nitrogen-containing compounds. Biology Bulletin Reviews 11:198–212.

28. Kubička J. 1966. Čtyři případy otravy ucháčem (Gyromitra). Česká Mykologie 20:178–181.

29. Lagrange E, Vernoux JPP, Reis J, Palmer V, Camu W, Spencer PSS. 2021. An amyotrophic lateral sclerosis hot spot in the French Alps associated with genotoxic fungi. Journal of the Neurological Sciences 427:1–8.

30. Larget B, Simon DL. 1999. Markov chain Monte Carlo algorithms for the Bayesian analysis of phylogenetic trees. Molecular Biology and Evolution 16:750–759.

31. Leathem AM, Dorran TJ. 2007. Poisoning due to raw *Gyromitra esculenta* (false morels) west of the Rockies. Canadian Journal of Emergency Medicine 9:127–130.

32. Li H-J, Zuo-Hong C, Cai Q, Zhou M-H, Chen G-J, Cheng-Ye S, Zhang H-S, Zhu-Liang Y. 2020. *Gyromitra venenata*, a new poisonous species discovered from China. Mycosystema 39:1706– 1718.

33. Liang Y-H, Eisenga BH, Trestrail JTI, Kuslikis B. 1998. Gyromitra mushroom species and their monomethylhydrazine content. In: Platform Session 1: Part 2, Journal of Toxicology: Clinical Toxicology. p. 527.

34. List PH, Luft P. 1968a. Gyromitrin, das gift der frühjahrslorchel. 16. Mitt. über pilzinhaltsstoffe [Gyromitrin, the poison of Gyromitra esculenta. 16. On the fungi contents]. Arch Pharm Ber Dtsch Pharm Ges 301:294–305.

35. List PH, Luft P. 1968b. Nachweis und gehaltsbestimmung von gyromitrin in frischen lorcheln. 19. Mitt. über pilzinhaltsstoffe [Detection and content determination of gyromitrin in fresh Gyromitra esculenta. 19. Mushroom contents]. Arch Pharm Ber Dtsch Pharm Ges 302:143–146.

36. Luo H, Hallen-Adams HE, Lüli Y, Sgambelluri RM, Li X, Smith M, Yang ZL, Martin FM. 2022. Genes and evolutionary fates of the amanitin biosynthesis pathway in poisonous mushrooms. PNAS 119:1–11.

37. Mäkinen SM, Kreula M, Kauppi M. 1977. Acute oral toxicity of ethylidene gyromitrin in rabbits, rats and chickens. Food and Cosmetics Toxicology 15:575–578.

38. Marjatta R, Pyysalo H. 1978. Occurrence of N-methyl-N-formylhydrazones in mycelia of *Gyromitra esculenta*. Zeitschrift Fur Naturforschung 33:472–474.

39. McIlvaine C, Macadam RK. 1902. One Thousand American Fungi. Indianapolis: Bowen-Merrill Company. 729 p.

40. Michelot D, Toth B. 1991. Poisoning by *Gyromitra esculenta*—a review. Journal of Applied Toxicology 11:235–243.

41. Miller AN, Dirks AC, Filippova N, Popov E, Methven AS. 2022. Studies in *Gyromitra* II: Cryptic speciation in the *Gyromitra gigas* species complex; rediscovery of *G. ussuriensis* and *G. americanigigas* sp. nov. Research Square 1–23.

42. Miller AN, Yoon A, Gulden G, Stensholt Ø, Vooren N Van, Ohenoja E, Methven AS. 2020. Studies in *Gyromitra* I: The *Gyromitra gigas* species complex. Mycological Progress 19:1459–1473.

43. Miller MA, Pfeiffer W, Schwartz T. 2011. The CIPRES science gateway: A community resource for phylogenetic analyses. In: Proceedings of the TeraGrid 2011 Conference: Extreme Digital Discovery, TG’11. p. 1–8.

44. Mohamed OG, Khalil ZG, Capon RJ. 2018. Prolinimines: N-amino-L-pro-methyl ester (hydrazine) Schiff bases from a fish gastrointestinal tract-derived fungus, Trichoderma sp. CMB-F563. Organic Letters 20:377–380.

45. Mohamed OG, Khalil ZG, Capon RJ. 2021. N-amino-L-proline methyl ester from an Australian fish gut-derived fungus: Challenging the distinction between natural product and artifact. Marine Drugs 19:1–14.

46. Moncalvo JM, Lutzoni FM, Rehner SA, Johnson J, Vilgalys R. 2000. Phylogenetic relationships of agaric fungi based on nuclear large subunit ribosomal DNA sequences. Systematic Biology 49:278–305.

47. Monteith WR. 2020. Draft Environmental Assessment for SpaceX Falcon Launches at Kennedy Space Center and Cape Canaveral Air Force Station. 110 p.

48. Morris JJ, Lenski RE, Zinser ER. 2012. The black queen hypothesis: Evolution of dependencies through adaptive gene loss. MBio 3:1–7.

49. Patron NJ, Waller RF, Cozijnsen AJ, Straney DC, Gardiner DM, Nierman WC, Howlett BJ. 2007. Origin and distribution of epipolythiodioxopiperazine (ETP) gene clusters in filamentous ascomycetes. BMC Evolutionary Biology 7:174–188.

50. Porter TM, Martin W, James TY, Longcore JE, Gleason FH, Adler PH, Letcher PM, Vilgalys R. 2011. Molecular phylogeny of the Blastocladiomycota (Fungi) based on nuclear ribosomal DNA. Fungal Biology 115:381–392.

51. Pyysalo H. 1975. Some new toxic compounds in false morels, *Gyromitra esculenta*. Naturwissenschaften 62:395.

52. Pyysalo H. 1976. Tests for gyromitrin, a poisonous compound in false morel *Gyromitra esculenta*. Zeitschrift Für Lebensmittel-Untersuchung Und -Forschung 160:325–330.

53. Pyysalo H, Niskanen A, von Wright A. 1978. Formation of toxic methylhydrazine during cooking of false morels, *Gyromitra esculenta*. Journal of Food Safety 1:295–299.

54. Pyysalo H, Niskanen AA. 1977. On the occurrence of N-methyl-N-formylhydrazones in fresh and processed false morel, *Gyromitra esculenta*. Journal of Agricultural and Food Chemistry 25:644– 647.

55. Rokas A, Mead ME, Steenwyk JL, Raja HA, Oberlies NH. 2020. Biosynthetic gene clusters and the evolution of fungal chemodiversity. Natural Product Reports 37:868–878.

56. Ronquist F, Teslenko M, Mark P Van Der, Ayres DL, Darling A, Höhna S, Larget B, Liu L, Suchard MA, Huelsenbeck JP. 2012. Mrbayes 3.2: Efficient bayesian phylogenetic inference and model choice across a large model space. Systematic Biology 61:539–542.

57. Roupas P, Keogh J, Noakes M, Margetts C, Taylor P. 2010. Mushrooms and agaritine: A mini-review. Journal of Functional Foods 2:91–98.

58. Sitta N, Davoli P, Floriani M, Suriano E. 2021. Guida ragionata alla commestibilità dei funghi. 180 p.

59. Spencer PS. 2020. Consumption of unregulated food items (false morels) and risk for neurodegenerative disease (amyotrophic lateral sclerosis). Health Risk Analysis 2020:93–98.

60. Spencer PS, Kisby GE. 2021. Role of hydrazine-related chemicals in cancer and neurodegenerative disease. Chemical Research in Toxicology 34:1953–1969.

61. Spencer PS, Palmer VS. 2021. Direct and indirect neurotoxic potential of metal/metalloids in plants and fungi used for food, dietary supplements, and herbal medicine. Toxics 9:57–69.

62. Stamatakis A. 2014. RAxML version 8: A tool for phylogenetic analysis and post-analysis of large phylogenies. Bioinformatics 30:1312–1313.

63. Svanberg I, Lindh H. 2019. Mushroom hunting and consumption in twenty-first century post-industrial Sweden. Journal of Ethnobiology and Ethnomedicine 15:1–23.

64. Talavera G, Castresana J. 2007. Improvement of phylogenies after removing divergent and ambiguously aligned blocks from protein sequence alignments. Systematic Biology 56:564–577.

65. Toth B, Nagel D. 1978. Tumors induced in mice by N-methyl-N-formylhydrazine of the false morel *Gyromitra esculenta*. Journal of the National Cancer Institute 60:201–204.

66. Toth B, Patil K. 1980. Carcinogenesis by a single dose of N-methyl-N-formylhydrazine. Journal of Toxicology and Environmental Health 6:577–584.

67. Tylutki EE. 1979. Mushrooms of Idaho and the Pacific Northwest: Vol. 1 Discomycetes. Moscow, Idaho: University Press of Idaho. 133 p.

68. Viernstein H, Jurenitsch J, Kubelka W. 1980. Vergleich des giftgehaltes der lorchelarten Gyromitra gigas, Gyromitra fastigiata und Gyromitra esculenta. Ernahrung/Nutrition 4:392– 395.

69. Wang X-CC, Zhuang W-YY. 2019. A three-locus phylogeny of *Gyromitra* (Discinaceae, Pezizales) and discovery of two cryptic species. Mycologia 111:69–77.

70. Weber NS. 1995. A Morel Hunter’s Companion: A Guide to True and False Morels. Holt, Michigan: Thunder Bay Press. 209 p.

71. White TJ, Bruns TD, Lee SB, Taylor JW, White TJ, Bruns TD, Lee SB, Taylor JW. 1990. Amplification and direct sequencing of fungal ribosomal RNA Genes for phylogenetics. In: PCR Protocols: A Guide to Methods and Applications. Academic Press, Inc. p. 315–322.

72. Wright A V., Niskanen A, Pyysalo H, Korpela H. 1978. The toxicity of some N-methyl-N-formylhydrazones from *Gyromitra esculenta* and related compounds in mouse and microbial tests. Toxicology and Applied Pharmacology 45:429–434.

73. Yancey CE, Smith DJ, Uyl PA Den, Mohamed OG, Yu F, Ruberg SA, Chaffin JD, Goodwin KD, Tripathi A, Sherman DH, Dick GJ. 2022. Metagenomic and metatranscriptomic insights into population diversity of *Microcystis* blooms: Spatial and temporal dynamics of mcy genotypes, including a partial operon that can be abundant and expressed. Applied and Environmental Microbiology 88:1–17.

