## Supplemental Information for "Distribution of the gyromitrin mycotoxin in the lorchel family assessed by a pre-column-derivatization and ultra high-performance liquid chromatography method"

**Supplemental Figures**

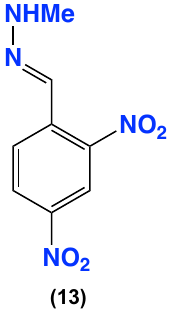

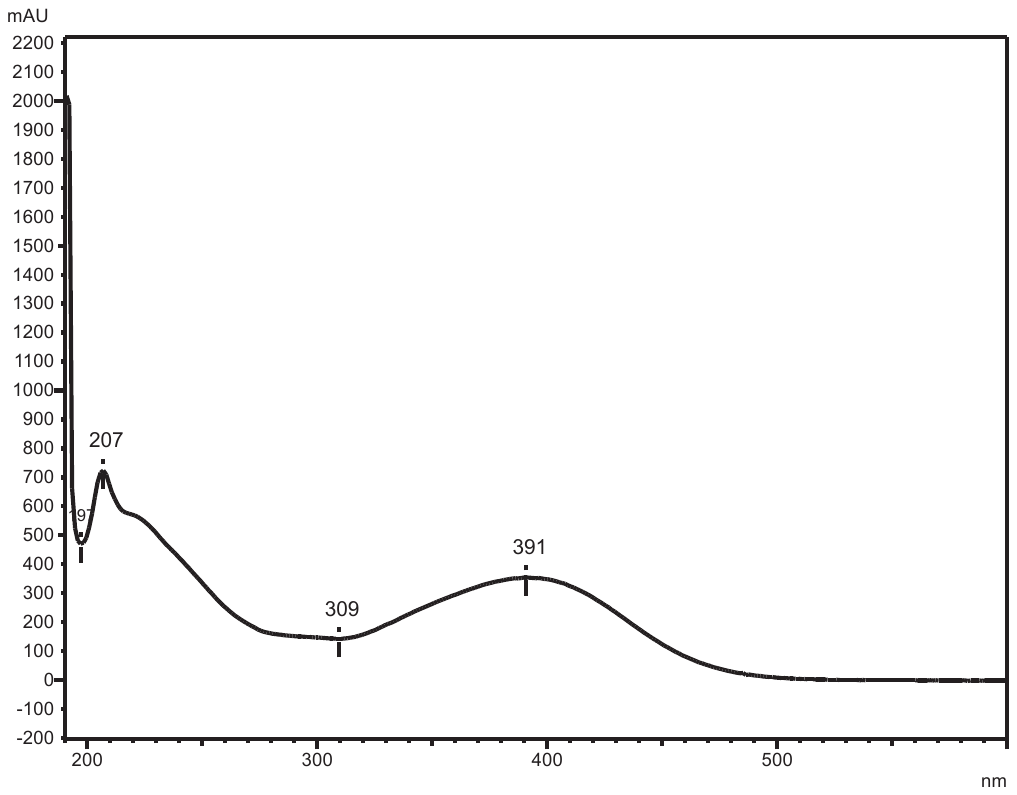

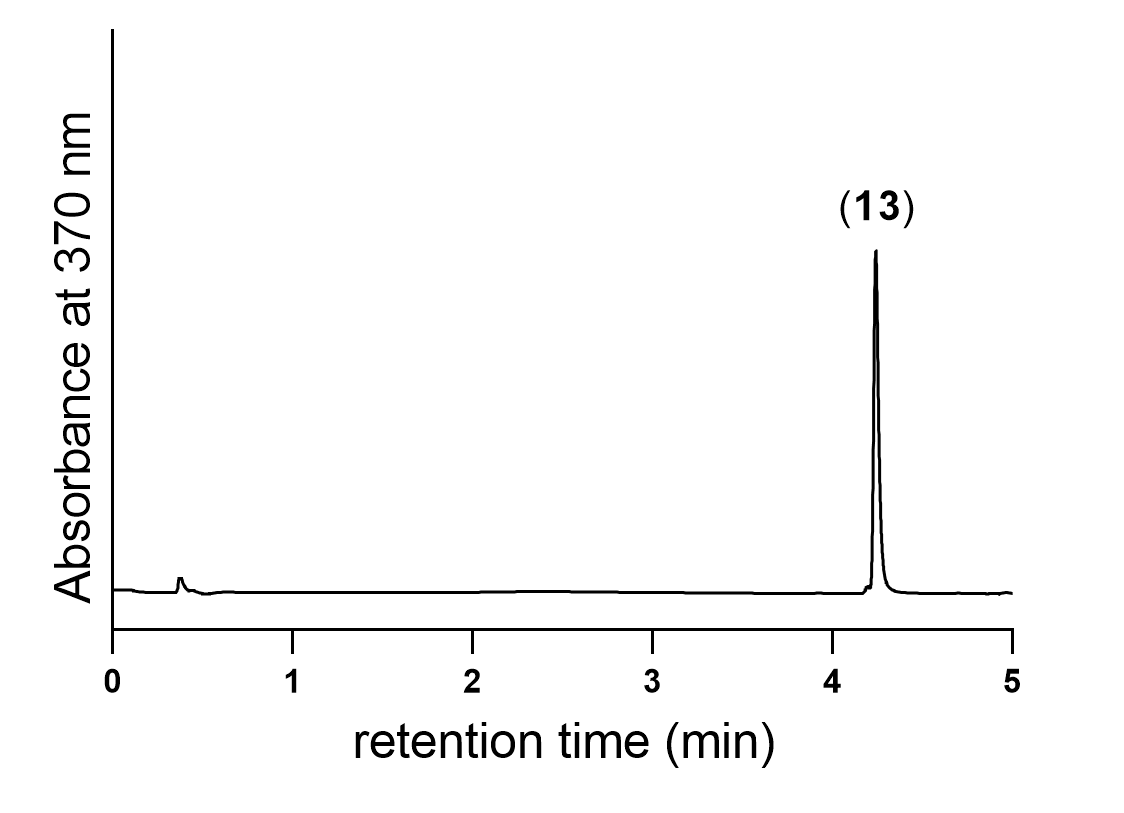

**Supplemental Figure 1.** UHPLC-DAD (370 nm) trace and UV-vis spectrum of synthetic monomethylhydrazine Schiff base (**13**).

| Concentration (mg/mL) | Peak area |
| --- | --- |
| 0.39 | 34001.5 |
| 0.78 | 67171 |
| 1.56 | 127711 |
| 3.12 | 250692 |
| 6.25 | 484315 |
| 12.5 | 966328 |
| 25 | 1810842 |
| 50 | 3434101 |

**Supplemental Figure 2.** Standard calibration curve of synthetic monomethylhydrazine Schiff base (**13**) at different concentrations (µg/mL) against UV peak area (370 nm).

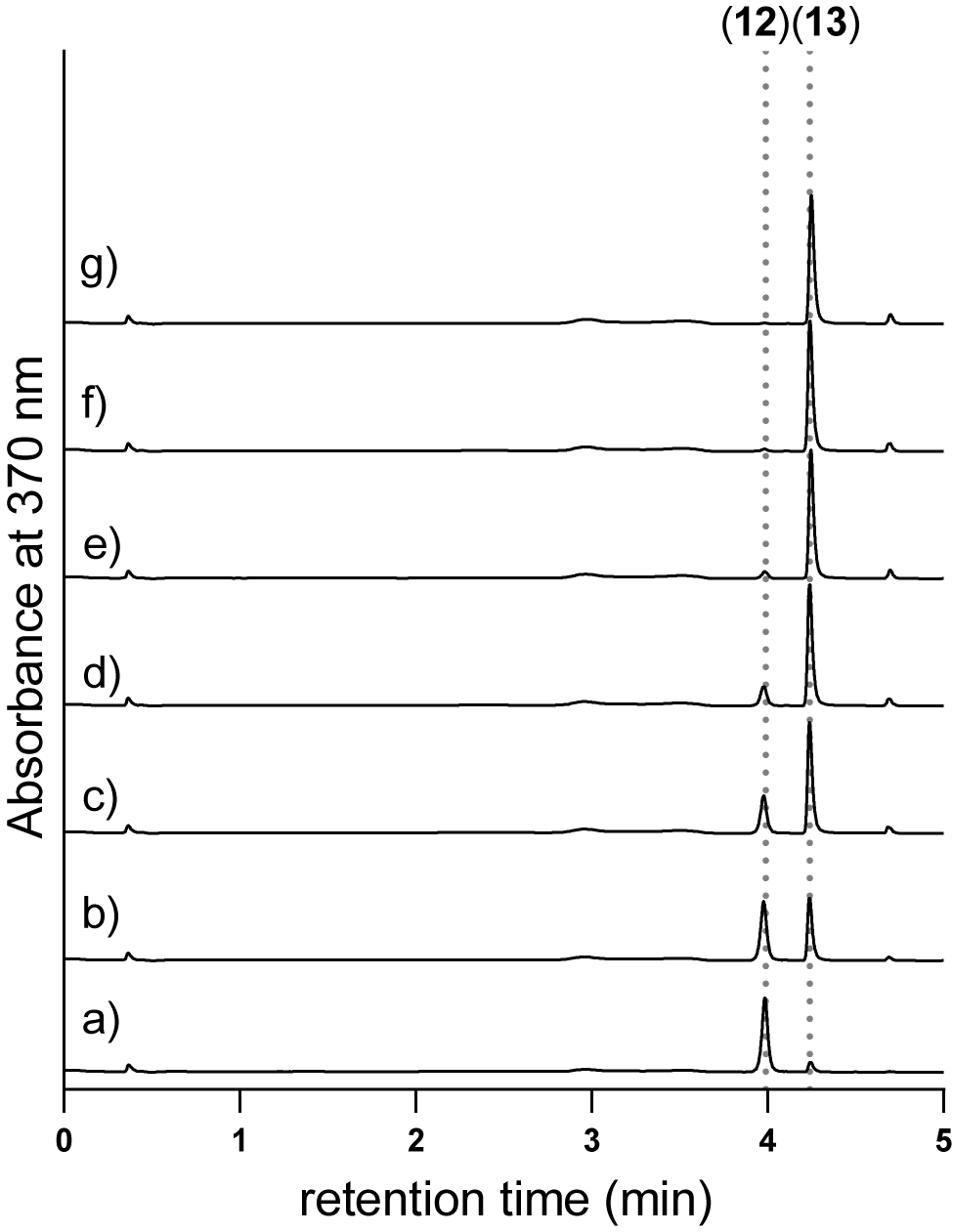

**Supplemental Figure 3.** UHPLC-DAD (370 nm) chromatograms of derivatization progress of gyromitrin standard (20 µg/mL) with 2,4-DNB at different time intervals. a) 0 h, b) 2 h, c) 5 h, d) 8 h, e) 13 h, f) 18 h, and g) 24 h.

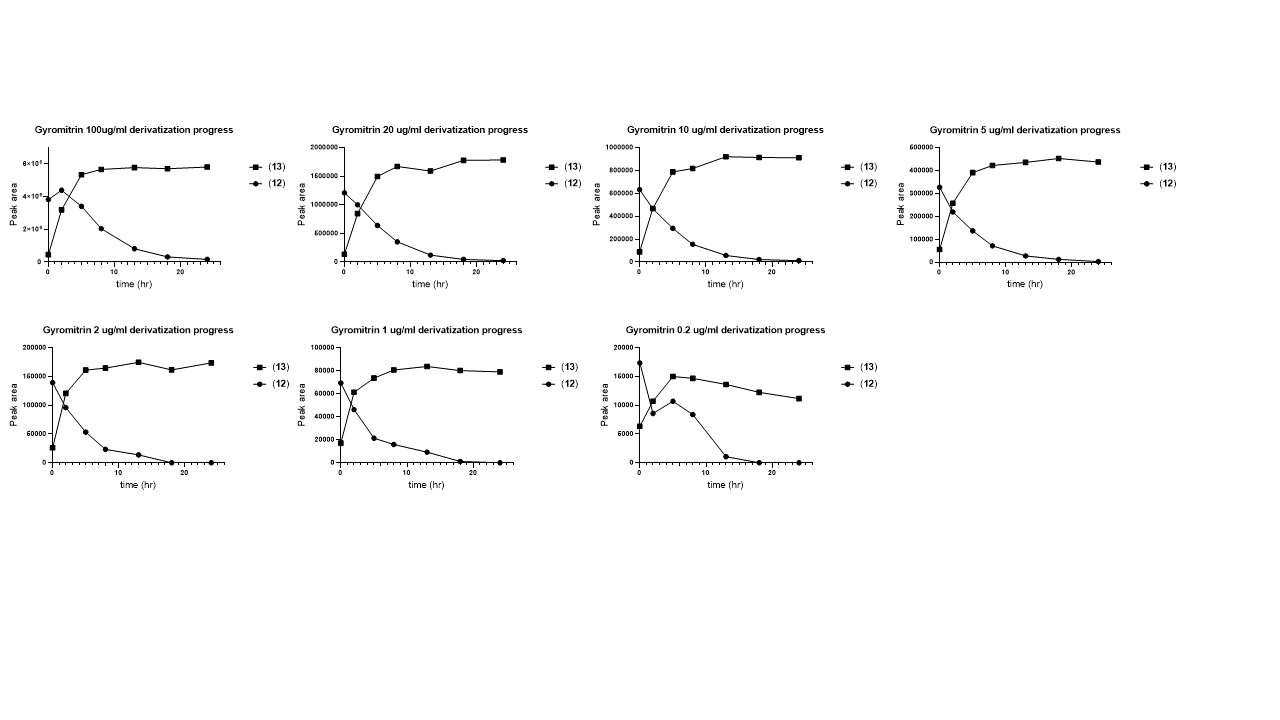

**Supplemental Figure 4.** Hydrolysis and derivatization progress of different concentrations of gyromitrin standard using 2,4-DNB over time.

| Concentration (mg/mL) | Peak area |
| --- | --- |
| 20 | 1775369 |
| 10 | 913033 |
| 5 | 451864 |
| 2 | 161334 |
| 1 | 80118 |

**Supplemental Figure 5.** Correlation between different concentrations of gyromitrin standard solution and UV peak area (370 nm) of derivatized product (**13**).

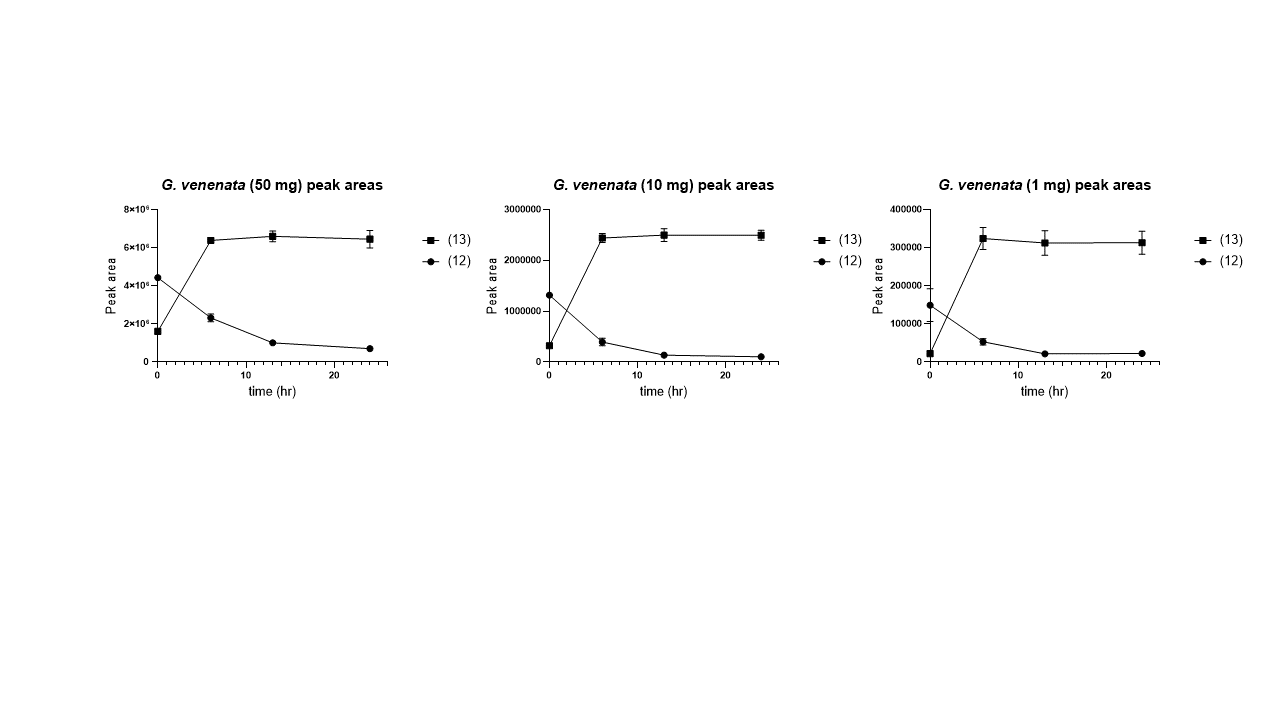

**Supplemental Figure 6.** Hydrolysis and derivatization progress of gyromitrin in different amounts of *Gyromitra venenata* (MICH 352032) over time using 2,4-DNB .

**Supplemental Tables**

**Supplemental Table 1.** Accession numbers of GenBank sequences used in the phylogenetic analyses.

| **Original Determination** | **Fungarium** | **ID (collector number, isolate, strain)** | **Location** | **ITS** | **LSU** | **Type** | **Reference(s)** |
| --- | --- | --- | --- | --- | --- | --- | --- |
| *Gyromitra accumbens* | O86249 | NA | Norway |  | KX008332 |  | Miller unpublished |
| *Gyromitra antarctica* | FLAS-F-63238 | MES-1298 | Argentina | MH930384 | MZ018861 |  | Mujic and Smith unpublished; Healy et al. 2022 |
| *Gyromitra anthracobia* | LIP0001407 | NA | Cyprus | MH014751 | MH014750 | HOLOTYPE | Crous et al. 2018 |
| *Gyromitra anthracobia* | NA | ML71422V2 | Cyprus | MH014752 | MH014749 |  | Crous et al. 2018 |
| *Gyromitra californica* | OSC69668 | Mike Roantree s.n. | USA-OR | EU837203 |  |  | Gordon unpublished |
| *Gyromitra caroliniana* | NY01797002 | KUO-04100601 | USA-IL |  | KC751528 |  | Methven et al. 2013 |
| *Gyromitra esculenta* | WSP72380 | Ge 8-1 | USA-ID | KM204673 | KM204699 |  | Carris et al. 2015 |
| *Gyromitra esculenta* | HMAS214602 | NV2015.04.01 | France | MG846990 | MG847002 | EPITYPE | Wang and Zhuang 2019 |
| *Gyromitra fastigiata* | HMAS254603 | NV2004.05.01 | France | MG846992 | MG847003 |  | Wang and Zhuang 2019 |
| *Gyromitra gigas* | TUR-A208088 | NA | Czech Republic | MH938663 | MH938309 | EPITYPE | Carbone et al. 2018 |
| *Gyromitra infula* | HMAS250470 | NA | China | MG846956 | MG847006 |  | Wang and Zhuang 2019 |
| *Gyromitra korfii* | CUP28997 | NA | USA-NC | MW075385 | MW078425 | PARATYPE | Miller et al. 2020 |
| *Gyromitra leucoxantha* | HMAS279665 | NV2017.06.10 | Spain | MG846991 | MG847020 |  | Wang and Zhuang 2019 |
| *Gyromitra melaleucoides* | MICH1455 | NSW 4520 | USA-CO |  | KC751517 |  | Methven et al. 2013 |
| *Gyromitra montana* | BPI 566707 | NA | USA-WY | MW077452 | MW077442 | ISOTYPE | Miller et al. 2020 |
| *Gyromitra olympiana* | FLAS-F-61941 | MES-3061 | USA-WA | MT373908 | MT350422 |  | Healy unpublished |
| *Gyromitra perlata* | HMAS254607 | NV2014.05.14 | France | MG846993 | MG847022 |  | Wang and Zhuang 2019 |
| *Gyromitra pseudogigas* | HMAS46539 | NA | China | MG846994 | MG847023 | HOLOTYPE | Wang and Zhuang 2019 |
| *Gyromitra slonevskii* | HAI-D077 | D-077 | Ukraine | JQ691488 |  |  | Barseghyan et al. 2012 |
| *Gyromitra sphaerospora* | HMAS73375 | NA | China | MG846997 | MG847027 |  | Wang and Zhuang 2019 |
| *Gyromitra spinosospora* | ZT-Myc-64133 | NA | Switzerland | MW489531 |  |  | Senn-Irlet et al. 2021 |
| *Gyromitra splendida* | TAAM046650 | KL398 | Estonia | KX185090 | KX185094 | HOLOTYPE | Miller et al. 2015 |
| *Gyromitra tasmanica* | PDD83761 | JAC9664 | New Zealand | MK432693 |  |  | Cooper et al. unpublished |
| *Gyromitra tianshanensis* | HMAS86057 | NA | China | MG846963 | MG847024 | HOLOTYPE | Wang and Zhuang 2019 |
| *Gyromitra venenata* | HKAS107323 | NA | China | MT424856 | MT421932 | PARATYPE | Li et al. 2020 |
| *Hydnotrya brunneospora* | HMAS97138 | NA | China | NR_161073 |  | HOLOTYPE | Xu et al. 2018 |
| *Hydnotrya cerebriformis* | NA | PR980813_IT7 | Germany | GQ140236 |  |  | Stielow et al. 2010 |
| *Hydnotrya cubispora* | K104976 | B.J. Lack s.n. | Wales | EU784273 |  |  | Brock et al. 2009 |

**Supplemental Table 2.** Peak area and recovery percentage of **13** in the pre-column derivatization (18 hours post incubation at 40 C) and UHPLC-DAD analysis (370 nm) of gyromitrin standard at various concentrations.

| Gyromitrin standard concentration (µg/mL) | Peak area | Recovery percentage |
| --- | --- | --- |
| 100 | 5704347 | 36.80 |
| 20 | 1775369 | 56.33 |
| 10 | 913033 | 56.63 |
| 5 | 451864 | 53.32 |
| 2 | 161334 | 38.90 |
| 1 | 80118 | 25.02 |

**Supplemental Table 3.** Metadata and results of gyromitrin tests conducted on samples for this study.

| **Species** | **Fungarium Accession Number** | **Collection/Isolate Number** | **Origin** | **Year** | **Analyzed** | **Gyromitrin** |
| --- | --- | --- | --- | --- | --- | --- |
| *Cudonia grisea* | NA | iNat #51411067 | OR | 2020 | dried, frozen | 0 |
| *Disciotis* cf. *venosa* | MICH 352035 | ACD0290 | MI | 2020 | dried, frozen, culture (PDA) | 0 |
| *Gyromitra ambigua* | MICH 352091 | ACD0408 | CA | 2021 | dried ×2, culture (PDA) | 0 |
| *Gyromitra americanigigas* | MICH 352014 | ACD0256 | MI | 2020 | dried, frozen | 0 |
| *Gyromitra americanigigas* | MICH 352016 | ACD0258 | MI | 2020 | dried, frozen | 0 |
| *Gyromitra ancilis* | MICH 352024 | ACD0260 | MI | 2020 | dried, frozen | 0 |
| *Gyromitra ancilis* | MICH 352026 | ACD0263 | MI | 2020 | dried, frozen | 0 |
| *Gyromitra ancilis* | MICH 352031 | ACD0285 | MI | 2020 | frozen | 0 |
| *Gyromitra antarctica* | FLAS-F-65994 | NA | Chile | 2019 | dried | 1 |
| *Gyromitra brunnea* | MICH 352048 | ACD0324 | KA | 2020 | dried, frozen | 0 |
| *Gyromitra brunnea* | MICH 352054 | ACD0332 | MI | 2020 | dried | 0 |
| *Gyromitra californica* | ARZ-M-AN 01351 | NA | MO | 1977 | dried | 0 |
| *Gyromitra californica* | MICH 352088 | ACD0394 | WA | 2020 | dried ×2, culture (PDA) | 0 |
| *Gyromitra caroliniana* | MICH 352092 | ACD0409 | MO | 2021 | dried ×2 | 0 |
| *Gyromitra esculenta* | CBS 101906 | NA | Netherlands | culture | culture (PDA, MEA ×2) | 1 (PDA), 0 (MEA) |
| *Gyromitra* aff. *infula* | Betsy Arnold culture collection | DC3764 | southwestern USA | culture | culture (PDA) | 0 |
| *Gyromitra* aff. *infula* | MICH 28519 | NA | AZ | 1990 | dried ×2 | 0 |
| *Gyromitra infula* | Betsy Arnold culture collection | SO0215 | Panama | culture | culture (PDA) | 0 |
| *Gyromitra infula* | Betsy Arnold culture collection | SO4633 | Chile | culture | culture (PDA) | 0 |
| *Gyromitra infula* | CBS 113691 | NA | Sweden | culture | culture (PDA, MEA) | 0 |
| *Gyromitra infula* | MICH 352086 | ACD0393 | WA | 2020 | dried | 0 |
| *Gyromitra infula* | MICH 352090 | ACD0396 | WA | 2020 | dried ×2, culture (PDA) | 0 |
| *Gyromitra korfii* | MICH 352062 | ACD0399 | IN | 2021 | dried | 0 |
| *Gyromitra leucoxantha* | MICH 25407 | NA | MI | 1984 | dried | 1 |
| *Gyromitra leucoxantha* | MICH 352087 | ACD0326 | NY | 2020 | dried ×3 | 1 |
| *Gyromitra melaleucoides* | MICH 1455 | NA | CO | 1983 | dried | 0 |
| *Gyromitra melaleucoides* | MICH 352039 | ACD0308 | OR | 2020 | dried | 0 |
| *Gyromitra montana* | MICH 352043 | ACD0318 | OR | 2020 | dried, frozen | 0 |
| *Gyromitra* aff. *olympiana* | AN 044228 | NA | AZ | 2017 | dried | 0 |
| *Gyromitra* aff. *olympiana* | MICH 4614 | NA | ID | 1972 | dried ×2 | 0 |
| *Gyromitra* sp. | MICH 28557 | NA | AZ | 1983 | dried ×2 | 0 |
| *Gyromitra* sp. | MICH 4601 | NA | ID | 1962 | dried | 0 |
| *Gyromitra sphaerospora* | Betsy Arnold culture collection | WB0618 | western Canada | culture | culture (PDA) | 0 |
| *Gyromitra sphaerospora* | MICH 25703 | NA | MI | 1970 | dried ×2 | 0 |
| *Gyromitra sphaerospora* | MICH 25704 | NA | MI | 1970 | dried | 0 |
| *Gyromitra splendida* | MICH 25554 | NA | MI | 1984 | dried | 1 |
| *Gyromitra splendida* | MICH 352089 | ACD0395 | WA | 2020 | dried ×2, culture (PDA) | 1 |
| *Gyromitra venenata* | Betsy Arnold culture collection | IL0706 | north-central USA | culture | culture (PDA) | 1 |
| *Gyromitra venenata* | MICH 1304 | NA | ID | 1972 | dried | 1 |
| *Gyromitra venenata* | MICH 25528 | NA | MI | 1966 | dried | 1 |
| *Gyromitra venenata* | MICH 25541 | NA | MI | 1945 | dried | 1 |
| *Gyromitra venenata* | MICH 25543 | NA | MI | 1979 | dried | 1 |
| *Gyromitra venenata* | MICH 25555 | NA | NH | 1883 | dried | 1 |
| *Gyromitra venenata* | MICH 25573 | NA | MI | 1976 | dried | 1 |
| *Gyromitra venenata* | MICH 25578 | NA | MI | 1982 | dried | 1 |
| *Gyromitra venenata* | MICH 352028 | ACD0282 | MI | 2020 | dried, frozen | 1 |
| *Gyromitra venenata* | MICH 352029 | ACD0283 | MI | 2020 | dried, frozen | 1 |
| *Gyromitra venenata* | MICH 352030 | ACD0284 | MI | 2020 | dried, frozen, culture (PDA) | 1 |
| *Gyromitra venenata* | MICH 352032 | ACD0286 | MI | 2020 | dried, frozen, culture (PDA) | 1 |
| *Gyromitra venenata* | MICH 352033 | ACD0287 | MI | 2020 | dried, culture (PDA) | 1 |
| *Gyromitra venenata* | MICH 352034 | ACD0288 | MI | 2020 | dried | 1 |
| *Gyromitra venenata* | MICH 39037 | NA | MI | 1995 | dried | 1 |
| *Gyromitra venenata* | MICH 39039 | NA | MI | 1998 | dried | 1 |
| *Gyromitra venenata* | MICH 68345 | NA | MI | 2001 | dried | 1 |
| *Gyromitra venenata* | MICH 68493 | NA | MI | 2003 | dried | 1 |
| *Gyromitra warnei* | MICH 352073 | ACD0420 | MI | 2021 | dried | 0 |
| *Hydnotrya cerebriformis* | MICH 67763 | NA | AZ | 1996 | dried ×2 | 0 |
| *Hydnotrya* cf. *cubispora* | AN 043557 | NA | AZ | 2015 | dried | 0 |
| *Hydnotrya cubispora* | FLAS-F-62913 | NA | Scotland | 2018 | dried | 0 |
| *Hydnotrya michaelis* | MICH 70028 | NA | UT | 1995 | dried | 0 |
| *Leotia lubrica* | MICH 352041 | ACD0314 | MI | 2020 | dried, frozen | 0 |
| *Morchella punctipes* | NA | iNat #46602042 | WI | 2020 | frozen | 0 |
| *Pachycudonia monticola* | MICH 352040 | ACD0312 | OR | 2020 | dried, frozen | 0 |
| *Scutellinia* sp. | MICH 352036 | ACD0295 | MI | 2020 | dried | 0 |
| *Sphaerosporella brunnea* | MICH 352042 | ACD0317 | Canada | 2020 | dried, frozen, culture (PDA) |  |
| *Urnula craterium* | NA | iNat #42365327 | WI | 2020 | frozen | 0 |
